# TGM6, a helminth secretory product, mimics TGF-β binding to TβRII to antagonize TGF-β signaling in fibroblasts

**DOI:** 10.1101/2023.12.22.573140

**Authors:** Stephen E. White, Tristin A. Schwartze, Ananya Mukundan, Christina Schoenherr, Shashi P. Singh, Maarten van Dinther, Kyle T. Cunningham, Madeleine P. J. White, Tiffany Campion, John Pritchard, Cynthia S. Hinck, Peter ten Dijke, Gareth Inman, Rick M. Maizels, Andrew P. Hinck

## Abstract

The murine helminth parasite *Heligmosomoides polygyrus* expresses a family of proteins structurally related to TGF-β Mimic 1 (TGM1), a secreted five domain protein that activates the TGF-β pathway and converts naïve T lymphocytes to immunosuppressive Tregs. TGM1 signals through the TGF-β type I and type II receptors, TβRI and TβRII, with domains 1-2 and 3 binding TβRI and TβRII, respectively, and domains 4-5 binding CD44, a co-receptor abundant on T cells. TGM6 is a homologue of TGM1 that is co-expressed with TGM1, but lacks domains 1 and 2. Herein, we show that TGM6 binds TβRII through domain 3, but does not bind TβRI, or other type I or type II receptors of the TGF-β family. In TGF-β reporter assays in fibroblasts, TGM6, but not truncated TGM6 lacking domains 4 and 5, potently inhibits TGF-β- and TGM1-induced signaling, consistent with its ability to bind TβRII but not TβRI or other receptors of the TGF-β family. However, TGM6 does not bind CD44 and is unable to inhibit TGF-β and TGM1 signaling in T cells. To understand how TGM6 binds TβRII, the X-ray crystal structure of the TGM6 domain 3 bound to TβRII was determined at 1.4 Å. This showed that TGM6 domain 3 binds TβRII through an interface remarkably similar to the TGF-β:TβRII interface. These results suggest that TGM6 has adapted its domain structure and sequence to mimic TGF-β binding to TβRII and function as a potent TGF-β and TGM1 antagonist in fibroblasts. The coexpression of TGM6, along with the immunosuppressive TGMs that activate the TGF-β pathway, may prevent tissue damage caused by the parasite as it progresses through its life cycle from the intestinal lumen to submucosal tissues and back again.

## INTRODUCTION

Helminths, which have co-evolved with their mammalian hosts over long evolutionary timescales, persist by secreting soluble factors that suppress key immune signaling pathways and modulate host immunity [1-5]. In recent studies, we showed that upon infection, the murine intestinal helminth *Heligmosomoides polygyrus* secretes a protein known as TGF-β mimic, or TGM, that binds directly to the host receptors to activate the TGF-β pathway [6]. Thus, like the native cytokine, this stimulates the expression of the key transcriptional regulator Foxp3 in naïve T cells, expanding the population of CD4^+^ CD25^+^ Foxp3^+^ regulatory T cells (Tregs) [7, 8]. The increased numbers of Tregs promote peripheral immune tolerance and are required for the persistence of *H. polygyrus* in its mammalian host [2, 9-11].

The three mammalian TGF-β isoforms, TGF-β1, -β2, and -β3, control and influence many pathways in cellular differentiation [12-14] and are required for mediating immune tolerance [8, 12, 15] and maintaining vital tissues, such as the heart [14, 16]. The knockout of endogenous TGF-β1 in mice is characterized by the development of multi-organ inflammatory disease and death after maternal TGF-β1 is depleted [12]. The dysregulation of TGF-β signaling has been shown to drive the pathogenesis of several human diseases, including inflammatory bowel disease [17], renal, pulmonary, and cardiac fibrosis [18, 19], and cancer [18, 20, 21].

TGF-β growth factors are comprised of two elongated cystine-knotted monomers held together by a single interchain disulfide bond [22]. The growth factors signal by assembling a heterotetrameric complex with two near autonomously signaling pairs of serine/threonine kinase receptors, known as the TGF-β type I and type II receptors, TβRI and TβRII [23-26]. This triggers a phosphorylation cascade, with constitutively active TβRII phosphorylating TβRI, and activated TβRI phosphorylating the downstream transcriptional effector molecules, SMAD2 and SMAD3 [27].

In contrast to mammalian TGF-β, the helminth TGM molecule is a disulfide-rich 422 amino acid protein with an N-terminal signal peptide followed by five homologous domains [6]. Each domain has approximately 85 to 90 amino acids with either two or three disulfide bonds. These domains bear no homology to TGF-β or other TGF-β family members; instead, the individual domains are distantly related to the complement control protein (CCP) or sushi domain family [6, 28]. There are at least nine homologs of TGM in *H. polygyrus* [29], which are numbered TGM2 through TGM10, with the founding member, TGM, being numbered TGM1. Among this family of proteins, six (TGM1 – 6) are expressed primarily in the adult stages of the parasite while the remaining four (TGM7 - 10) are expressed in the larval stages [29, 30]. Domains 1, 2, and 3 (D1, D2, and D3, respectively) of TGM1 are necessary and sufficient for SMAD-dependent signaling in reporter cells [29], with D1 and D2 binding TβRI with a K_D_ of 30 nM and D3 binding TβRII with a K_D_ of 1.2 µM [28] (**Fig. 1A**). This combination of receptor pairings mimics the signaling complex created by TGF-β, though monomeric TGM1 binds and assembles a TβRII:TβRI heterodimer, while homodimeric TGF-β binds and assembles a TβRII:TβRI heterotetramer. Domains 4 and 5 (D4 and D5) bind CD44, a cell surface receptor that is abundant on T cells, which both targets and potentiates cellular responsiveness to TGM1 [31] (**Fig. 1A**).

**Figure 1.**
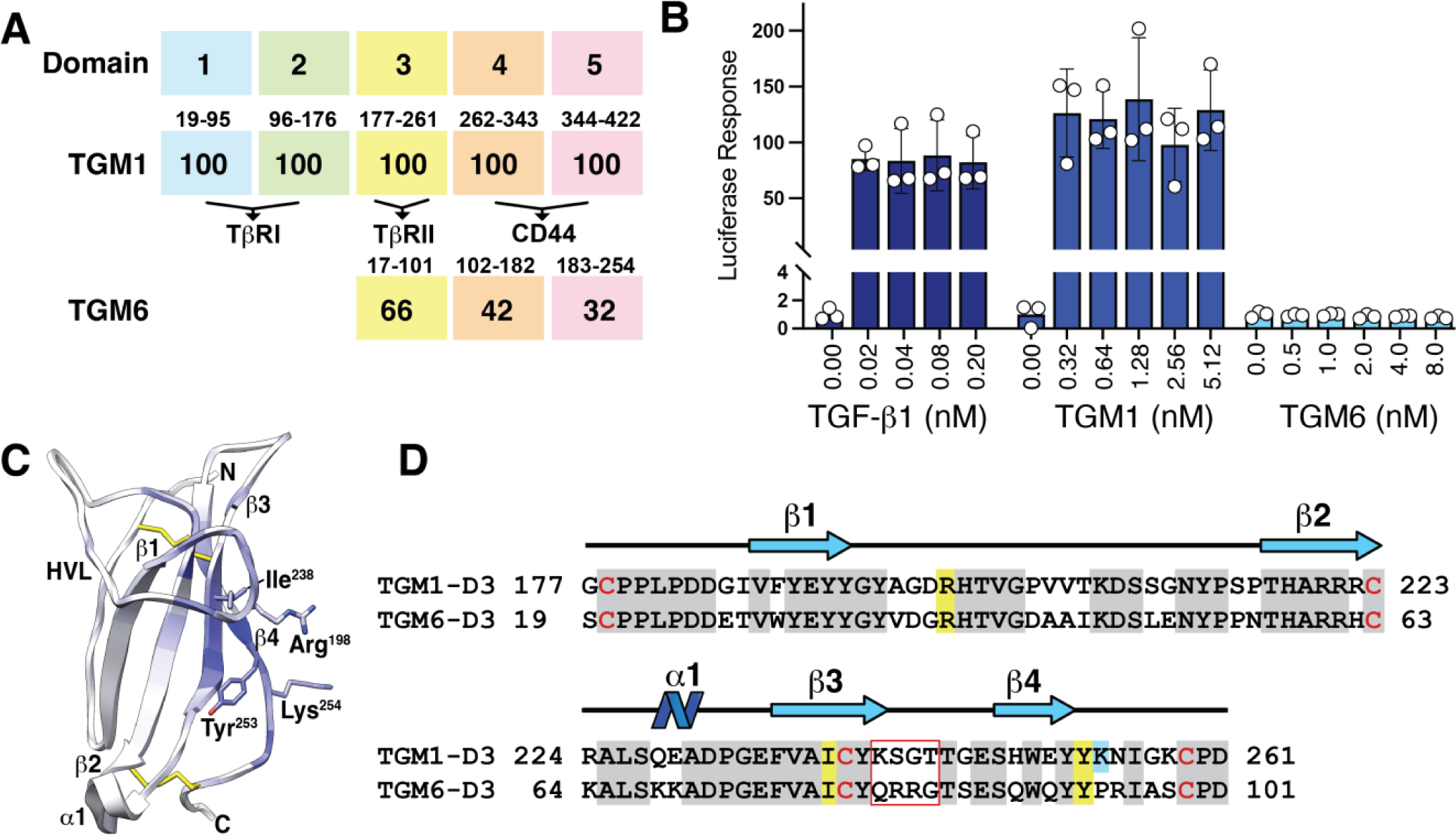
TGM6 domain structure, similarity to TGM1, and signaling activity. **A.** Comparison of the domain structure and amino sequence identity of TGM6 relative to TGM1 (large numbers in shaded boxes correspoding to each domain). Residue numbering is shown above the shaded boxes correspoding to each domain. **B.** TGF-β1-, TGM1- and TGM6-simulated luciferase reporter activity in NIH-3T3 fibroblasts. **C.** Structure of TGM1-D3 (PDB 7SXB) shaded according to chemical shift perturbations induced by binding of TβRII [28]. The sidechains of Arg^198^, Ile^238^, Tyr^253^, and Lys^254^ shown to be critical for binding are displayed [28]. **D.** Sequence alignment of TGM1-D3 and TGM6-D3. Conserved residues shown to be essential for binding TβRII by TGM1-D3 are highlighted in yellow (non-conserved are highlighted in cyan); all other conserved residues are shaded grey. Residues highlighted by red box are proposed to underlie the differential affinity of TGM6-D3 and TGM1-D3 for TβRII (see **Results** and Fig. 8). Note, residues numbers of TGM1-D3 and TGM6-D3 differ by 160 due the presence of D1-D2 only in TGM1. Figure shown in panel **C** adapted from Mukundan, et. al (2022), *J. Biol. Chem*, 298, 101994.

Among the TGMs expressed during the adult stages of the parasite, TGM6 is unique in that it lacks D1 and D2 (**Fig. 1A**) [6]. While TGM6 D3, D4, and D5 share 66%, 42%, and 32% identity with TGM1’s D3, D4, and D5, it does not signal in TGF-β functional assays, unlike TGM1, TGM2, and TGM3 which signal in both fibroblasts and T cells, or TGM4 which signals in T cells and myeloid cells [29, 32]. In this work, we show that TGM6 binds TβRII and that it does so through D3, but it does not bind TβRI or other type I and type II receptors of the TGF-β family. In TGF-β reporter assays in fibroblasts, TGM6 potently inhibits TGF-β- and TGM1-induced signaling, consistent with its receptor binding profile, including its ability to compete with TGF-β for binding TβRII. However, TGM6 does not inhibit TGF-β or TGM1 signaling in T cells, consistent with its divergent D4 and D5 and inability to bind CD44. We also present the crystal structure of the TGM6-D3:TβRII binary complex, in which TGM6-D3 is shown to fully mimic binding of mammalian TGF-β to TβRII, presenting a similar convex surface with a hydrophobic interior and charged residues on the periphery and engaging nearly the same set of residues. These results suggest that TGM6 has adapted its domain structure and sequence to mimic binding of mammalian TGF-β to TβRII and to antagonize TGF-β and TGM signaling in fibroblasts – but to do so without interfering with essential immune-suppressive signaling of TGM agonists in T cells.

## RESULTS

### TGM6 lacks signaling activity, but selectively and specifically binds TβRII

In contrast to mammalian-produced TGM1, TGM2, and TGM3, mammalian-produced TGM6 was previously shown to lack detectable signaling activity with the MFB-F11 reporter cell line, which is based on mouse embryonic fibroblasts and is specific for ligands that lead to activation of SMAD2/3 through its stably-transfected CAGA promoter element [33]. In consideration of the reported high specificity of the MFB-F11 reporter for signaling induced by TGF-βs, but not activins which also activate the SMAD2/3 branch of the TGF-β pathway, we used another murine reporter cell line, NIH-3T3 fibroblasts, also with a stably transfected CAGA promoter element, which are responsive to both TGF-βs and activins [34]. However, in accord with the previous MFB-F11 assay results, when tested at comparable concentrations, TGM6 was unable to activate the NIH-3T3 reporter above baseline, while both TGF-β1 and TGM1 robustly activated, with both nearly saturating the reporter even at the lowest concentrations tested (**Fig. 1B**).

In previous studies, we determined the structure of refolded bacterially expressed TGM1-D3 and showed that it adopted the overall fold of a CCP domain, but was expanded to open a potential interaction surface comprised of several residues near the C-terminus, as well as the tip of the long structurally ordered hypervariable loop (HVL), which wraps around the domain and extends toward β4, the C-terminal β-strand (**Fig. 1C**) [28]. Through NMR chemical shift perturbation mapping and site-directed mutagenesis and binding studies with TβRII, we showed TGM1-D3 contacts the same edge β-strand of TβRII (β4) as TGF-β; moreover, we identified three residues near the C-terminus of TGM1-D3, Ile^238^, Tyr^253^, and Lys^254^, and one residue on the tip of the hypervariable loop (HVL), Arg^198^, which when mutated led to a greater than 20-fold reduction in binding affinity (**Fig 1C**). Three of these four residues are identical in TGM6-D3, and the fourth, Lys^254^, has an arginine at an adjacent position in TGM6, suggesting that TGM6 might also bind the edge β-strand of TβRII through a similar surface (**Fig. 1D**).

Isothermal titration calorimetry (ITC) was used to determine if, in the absence of signaling activity, TGM6 might nonetheless bind TβRII. In titrations in which TβRII was titrated into mammalian produced TGM6, strong exothermic responses were observed, which after integration and fitting to a standard binding isotherm, yielded a K_D_ of 220 ± 100 nM (**Fig. 2A, Table S1**). To determine if TGM6-D3 holds the full capacity for binding TβRII, we produced ^15^N-TGM6-D3 in bacteria and showed using NMR that the refolded protein was natively folded (**Fig. S1A**). Titration of ^15^N-labeled TGM6-D3 with unlabeled TβRII led to perturbations and slow-exchange binding of more than half the amide signals of TGM6-D3, indicative of specific high affinity binding (**Fig. 2B**). In ITC measurements with TGM6-D3 and TβRII, strong exothermic binding was observed, which after integration and fitting to a standard binding isotherm, yielded a K_D_ of 440 ± 80 nM (**Fig. 2C**, **Table S1**). The difference between the K_D_ for TGM6 and TGM6-D3 for binding TβRII is not statistically significant (unpaired t-test p-value = 0.1826; assuming n = 2 for TGM6 and n = 3 for TGM6-D3 due to replicate experiment count). Thus, like TGM1, domain 3 holds the full capacity for binding TβRII, yet unlike TGM1, the affinity of TGM6 for TβRII is 4- to 5-times greater [28].

**Figure 2.**
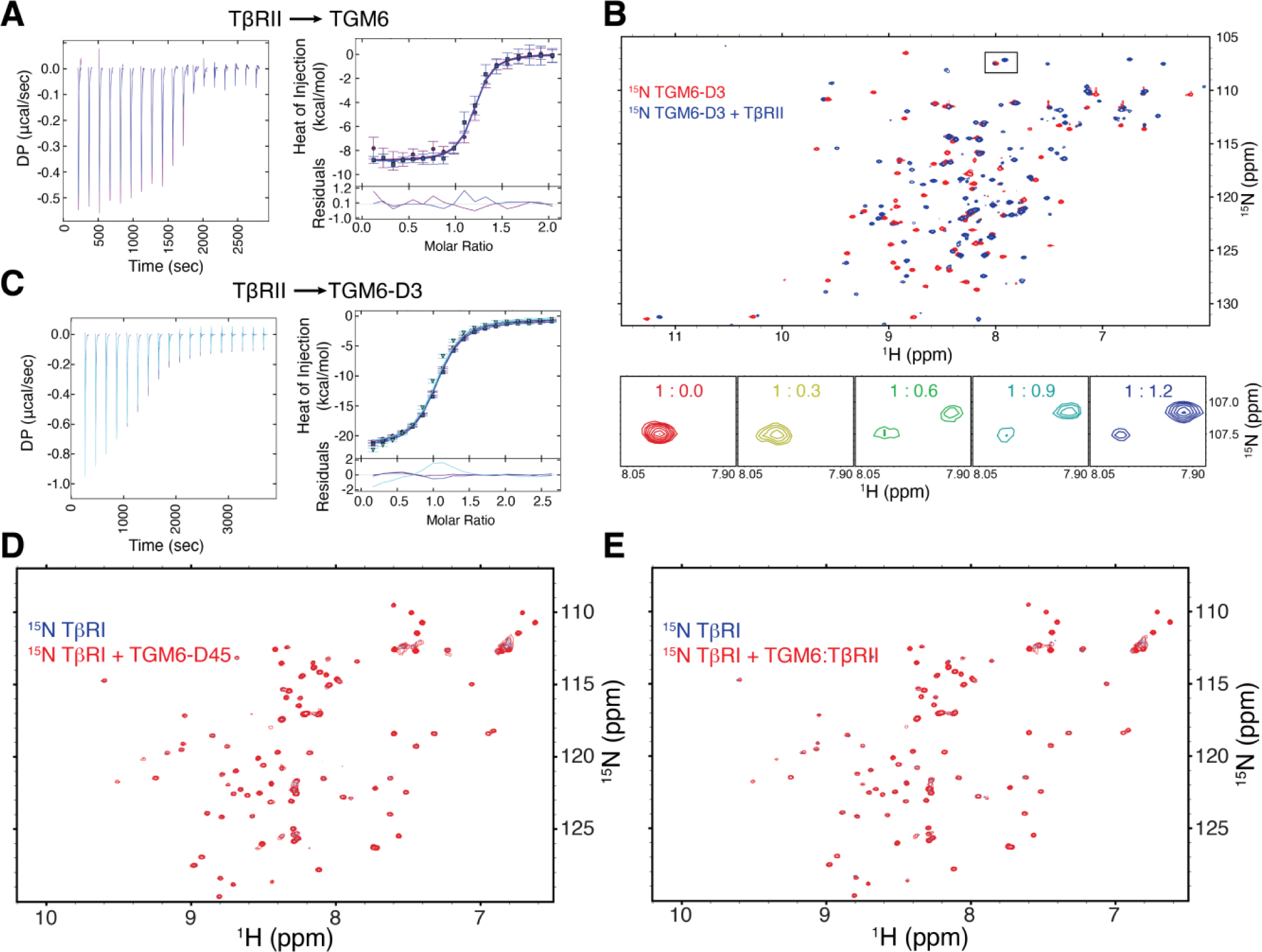
TGM6 binds TβRII through D3. **A,C.** ITC thermograms (left) obtained upon injection of TβRII into TGM6 (**A**) or TGM6-D3 (**C)**. Thermograms are overlaid as two (**A**) or three (**C**) replicates. Integrated heats are shown to the right of the thermograms with the residuals as a function of the molar ratio of TGM6:TβRII (**A**) or TGM6-D3:TβRII (**C**). The data points correspond to the integrated heats, and the colored lines correspond to a global fit of the data to a 1:1 binding model. **B.** ^1^H-^15^N HSQC spectra of ^15^N TGM6-D3 alone (red) overlaid onto the spectrum of the same sample containing a 1.2-fold molar excess of unlabeled TβRII (blue). Shown below is an expansion of the boxed region with all titration points labeled as the molar ratio of ^15^N TGM6-D3:TβRII. **D-E**. ^1^H-^15^N HSQC spectra of ^15^N TβRI alone (red) overlaid onto the spectrum of the same sample containing a 1.2-fold molar excess of unlabeled TGM6-D45 (blue) or TGM6:TβRII complex (blue) (**D** and **E**, respectively).

In the absence of TGM6 signaling activity in the MFB-F11 reporter line, we expected that TGM6 bound TβRI weakly, or not at all. To test this, we recorded NMR spectra of ^15^N-labeled TβRI alone and with 1.1 molar equivalents of unlabeled TGM6-D45 added (**Fig. 2D**). The addition of unlabeled TGM6 D45 led to no significant shifts in the signals of TβRI, even though the concentration was 100 μM. This indicates that TGM6 D45 does not directly bind TβRI, even with moderate affinity. To test the possibility that binding of TβRI is potentiated by TβRII, we re-recorded the spectra of ^15^N-TβRI, but with addition of 1.1 equivalents of unlabeled TGM6:TβRII complex, rather than TGM6-D45 alone (**Fig. 2E**). This also led to no significant shifts in the signals of TβRI, indicating that the TGM6:TβRII complex also does not bind TβRI. The absence of shifts of ^15^N-TβRI is not likely due to non-native folding of either TβRI or TGM6 D45, which were produced in *E. coli* and refolded, as the spectra of each are well-dispersed (**Fig. 2D-E, Fig. S1B**).

To test possible binding of TGM6-D3 by other type II receptors of the TGF-β family, we used ITC to determine whether TGM6-D3 binds the BMP and activin type II receptors ActRII, ActRIIB, and BMPRII (**Fig. S2, Table S2**). In this experiment, each type II receptor was titrated into TGM6-D3 or buffer alone and in each case, there was no detectable binding. To test possible binding of TGM6 by other type I receptors of the TGF-β family, we prepared ^15^N-labeled BMP and activin type I receptors, ALK1, ALK2, ALK3, and ALK4, and recorded spectra with 1.1 equivalents of unlabeled TGM6-D45 (**Fig. S3**) or TGM6:TβRII complex (**Fig. S4**) added, but like ^15^N-TβRI, no shifts were observed. The native folding of the type I and type II receptors is demonstrated by the chemical shift dispersion of their 2D ^1^H-^15^N (**Fig. S3, S4**) or 1D ^1^H spectra (**Fig. S5**), ruling out the possibility that misfolding of these disulfide-rich receptors is responsible for the lack of binding. Thus, TGM6 also does not bind the BMP and activin type I and type II receptors, either directly or in the case of type I receptors, by the TGM6:TβRII complex.

### TGM6-D3 competes for the TGF-β binding site on TβRII

TGM6-D3 binds TβRII and the key residues for binding in TGM1-D3 are conserved in TGM6-D3. Thus, it was hypothesized that TGM6-D3 would compete with TGF-β for binding TβRII, in a manner similar to TGM1-D3 (45). To test this, an ITC competition experiment was performed in which TβRII was titrated into the engineered TGF-β monomer, mmTGF-β2-7M2R [35], either alone or in the presence of 6 µM TGM6-D3 (**Fig. 3A**). The engineered TGF-β monomer was used for these experiments, rather than a native TGF-β dimer, due to its much higher solubility, yet unchanged TβRII binding affinity compared to the native dimer.

**Figure 3.**
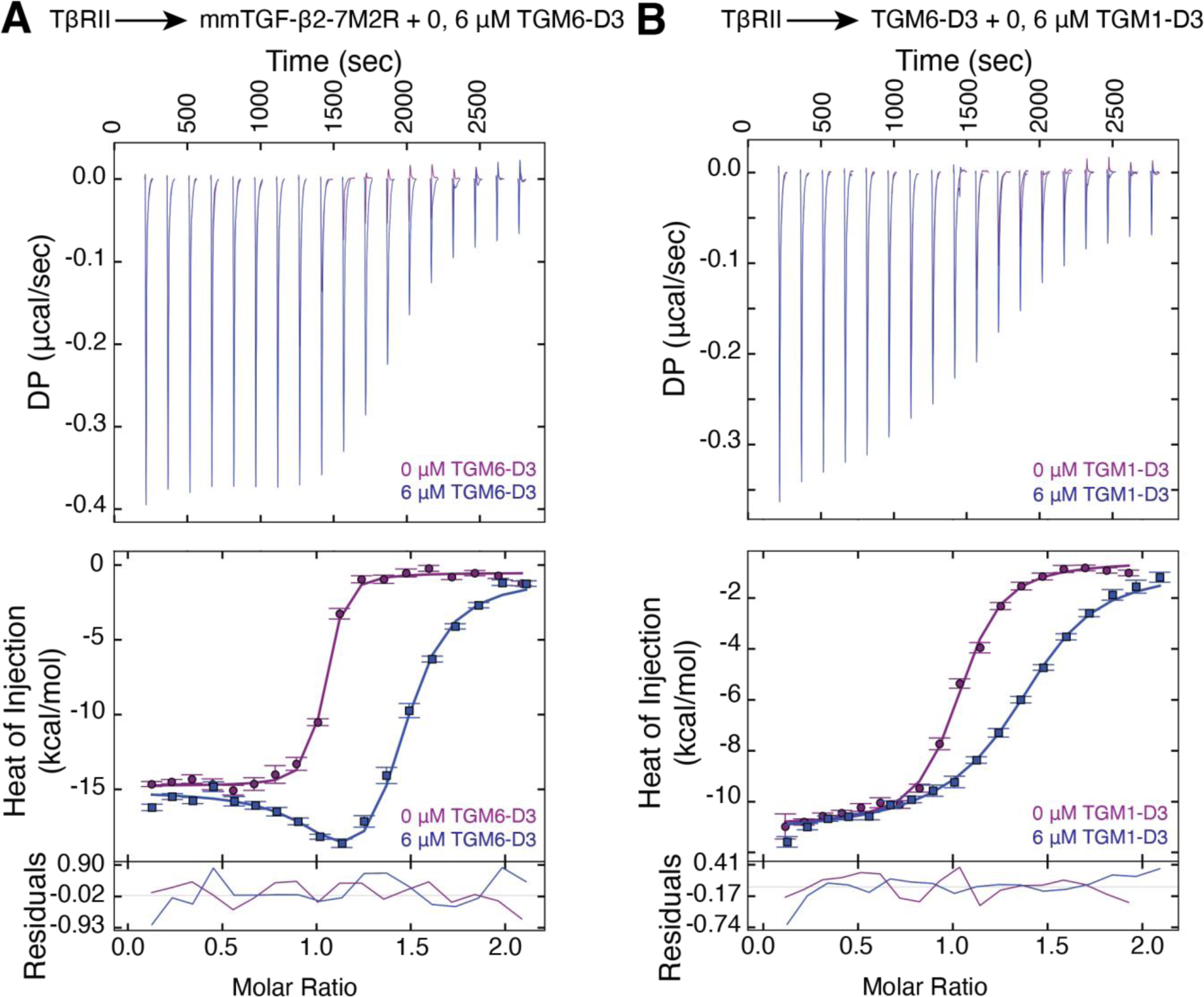
TGM6-D3, and TGM1-D3, compete with TGF-β for binding TβRII. (A) Titration of TβRII into mmTGF-β2-7M2R in the absence (purple) or presence (blue) of 6 μM TGM6-D3 as the lower-affinity binder. (B) Titration of TβRII into TGM6-D3 in the absence (purple) or presence (blue) of 6 μM TGM1-D3 as the lower-affinity binder. Each panel includes the thermograms (top), fitted isotherms (middle) and fitting residuals (bottom) for the associated titrations.

The addition of TGM6-D3 increased the extent of curvature in the binding isotherms and reduced the overall enthalpy of the reaction, consistent with the behavior expected for competitive binding (48). To quantify this interaction, the integrated heats from the two experiments were globally fit to a simple competitive binding model. The K_D_ and enthalpy for the lower-affinity TGM6-D3:TβRII interaction were held constant, and the binding parameters for the high-affinity mmTGF-β2-7M2R:TβRII interaction in the absence of the competitor were derived (**Table S3**). The K_D_ for the high-affinity mmTGF-β2-7M2R:TβRII was calculated to be 19 nM (±1σ confidence interval: 7.8 nM – 40 nM), which is in rough agreement with the previously determined K_D_ between mmTGF-β2-7M and TβRII of ca. 50 nM (49). Therefore, TGM6-D3 and TGF-β compete for the same binding site on TβRII. TGM6-D3 and TGM1-D3 are also expected to compete with one another for binding TβRII, based on their competitive binding with TGF-β. This was confirmed by performing a similar competition binding experiment, with TβRII titrated into TGM6-D3 either alone or in the presence of 6 µM TGM1-D3 (**Fig. 3B, Table S3**).

### TGM6 antagonizes TGF-β signaling in fibroblasts, but not T cells

TGM6 binds TβRII, but it does not bind TβRI, or any of the other BMP and activin type I and type II receptors tested, thus TGM6 might function as a TGF-β or TGM1 antagonist by occupying cell surface TβRII. To test this, NIH-3T3 fibroblast reporter cells were incubated with increasing concentrations of TGM6 prior to stimulation with TGF-β1 or TGM1 (**Fig. 4A**). The addition of TGM6 led to a dose-dependent decrease in signaling, with an IC_50_ of approximately 0.05 nM for inhibition of both TGF-β1 and TGM1. The assay was repeated using the MFB-F11 reporter fibroblasts and TGM6 similarly led to a dose-dependent decrease in signaling, with an IC_50_ of approximately 0.2 nM for inhibition of both TGF-β1 and TGM1 (**Fig. 4B**). To further characterize the inhibitory activity of TGM6, its ability to antagonize the conversion of murine splenic T cells to Foxp3^+^ Tregs by either TGF-β1 or TGM1 was measured, but no inhibition was observed, even at concentrations that nearly fully inhibited TGF-β1 or TGM1 signaling in the NIH-3T3 or MFB-F11 reporter cells (**Fig. 4C**).

**Figure 4.**
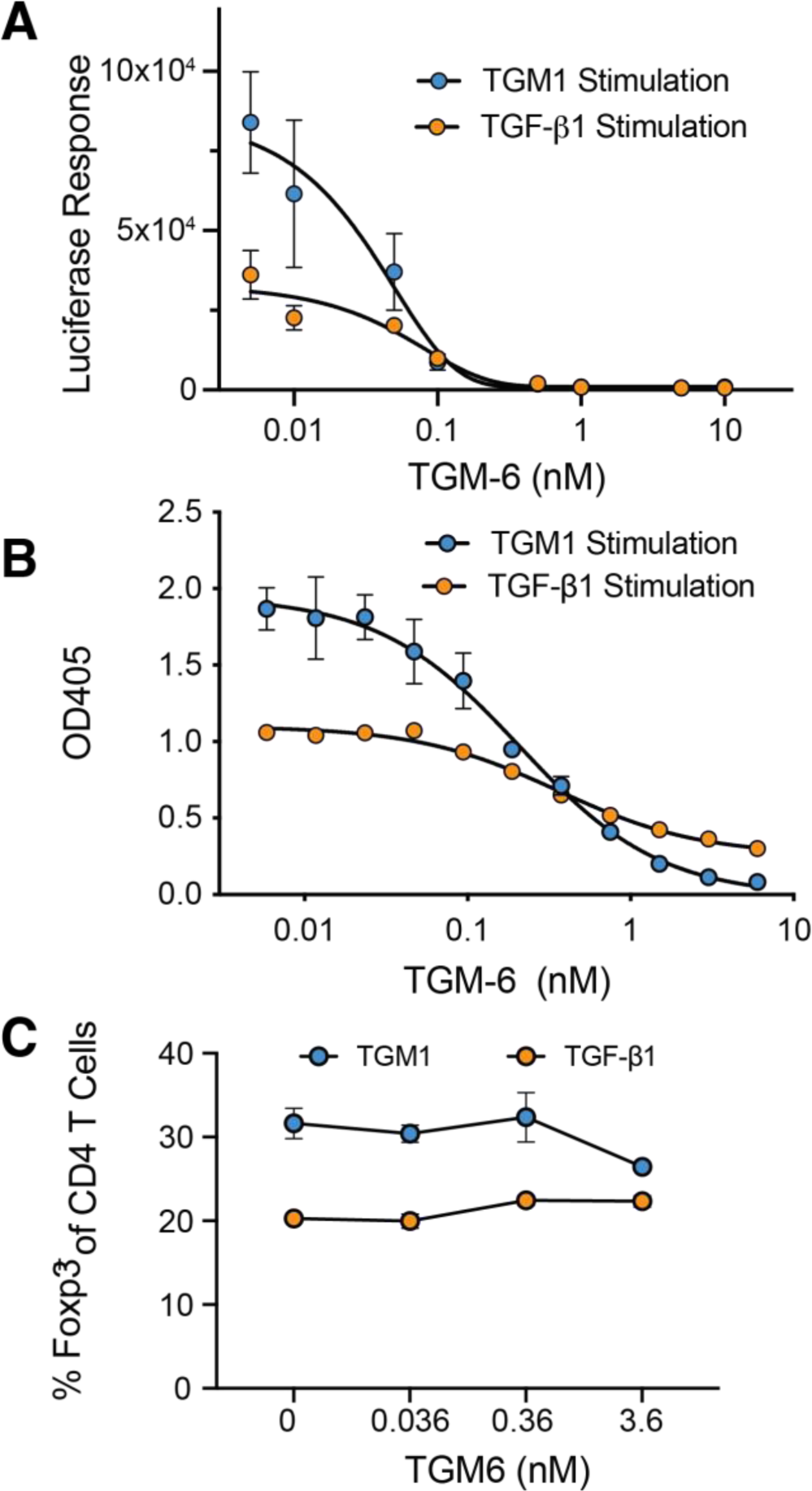
TGM6 is a potent inhibitor of TGF-β and TGM1 signaling in fibroblasts, but not T cells. **A-B.** Inhibition of SMAD2/3 CAGA reporter stimulated by TGF-β1 or TGM1 in NIH-3T3 (**A**) or MFB-F11 (**B**) fibroblasts by increasing concentrations of TGM6. Smooth black lines correspond to the fit of the data to a dose-dependent inhibition of TGF-β1 or TGM1 signaling by TGM6. **C.** Inhibition of the TGF-β1 or TGM1 induction of the Foxp3 transcription factor in murine splenic CD4+ T cells by increasing concentrations of TGM6.

To test if TGM6 could serve as either an agonist or antagonist of BMP signaling, we used NIH-3T3 fibroblasts stably transfected with a BMP responsive element (BRE) coupled to a fluorescent (mCherry) reporter and treated the cells with either BMP2, BMP6, or BMP7 alone or with 3.56 nM TGM6 added. The BMPs stimulated the reporter, but TGM6 was neither capable of stimulating the reporter itself nor inhibiting reporter activity stimulated by the BMPs (**Fig. S6A**). We also tested TGM6 for its ability to inhibit activin signaling using the NIH-3T3 fibroblasts stably transfected with a SMAD2/3 CAGA reporter element coupled to a fluorescent (GFP) reporter but observed that TGM6 was incapable of inhibiting activation of the reporter by activin A (ActA) (**Fig. S6B**).

The finding that TGM6 inhibits TGF-β1 and TGM1 signaling in fibroblasts, but not splenic T cells, and that its inhibitory concentration in both NIH-3T3 and MFB-F11 fibroblasts is several thousand-fold lower than its affinity for TβRII (ca. 0.1 nM vs. ca. 320 nM), suggests that its activity may be enhanced by a co-receptor, expressed by fibroblasts but not T cells, that is specifically recognized and bound by domains 4 and 5 (D45). To investigate this, the NIH-3T3 and MFB-F11 reporter assays were repeated but using TGM6-D3 for inhibition instead of full-length TGM6 (**Fig. 5A, B**). This resulted in no inhibition below 1000 nM in either cell line or only minor inhibition in the MFB-F11 cell line at concentrations higher than this, suggesting that D45 plays a critical role in the inhibition. To determine if physical attachment of D3 to D45 was required for inhibition, we repeated the inhibition assays in NIH-3T3 fibroblasts with treatment with D3, D45, or the combination of D3 and D45 (**Fig. 5C**). However, in contrast to the full-length protein at a concentration of 3.6 nM which completely inhibited signaling induced by TGF-β, there was no inhibition by the individual domains, or the combination at the same concentrations, suggesting that physical attachment of the domains is required for inhibition.

**Figure 5.**
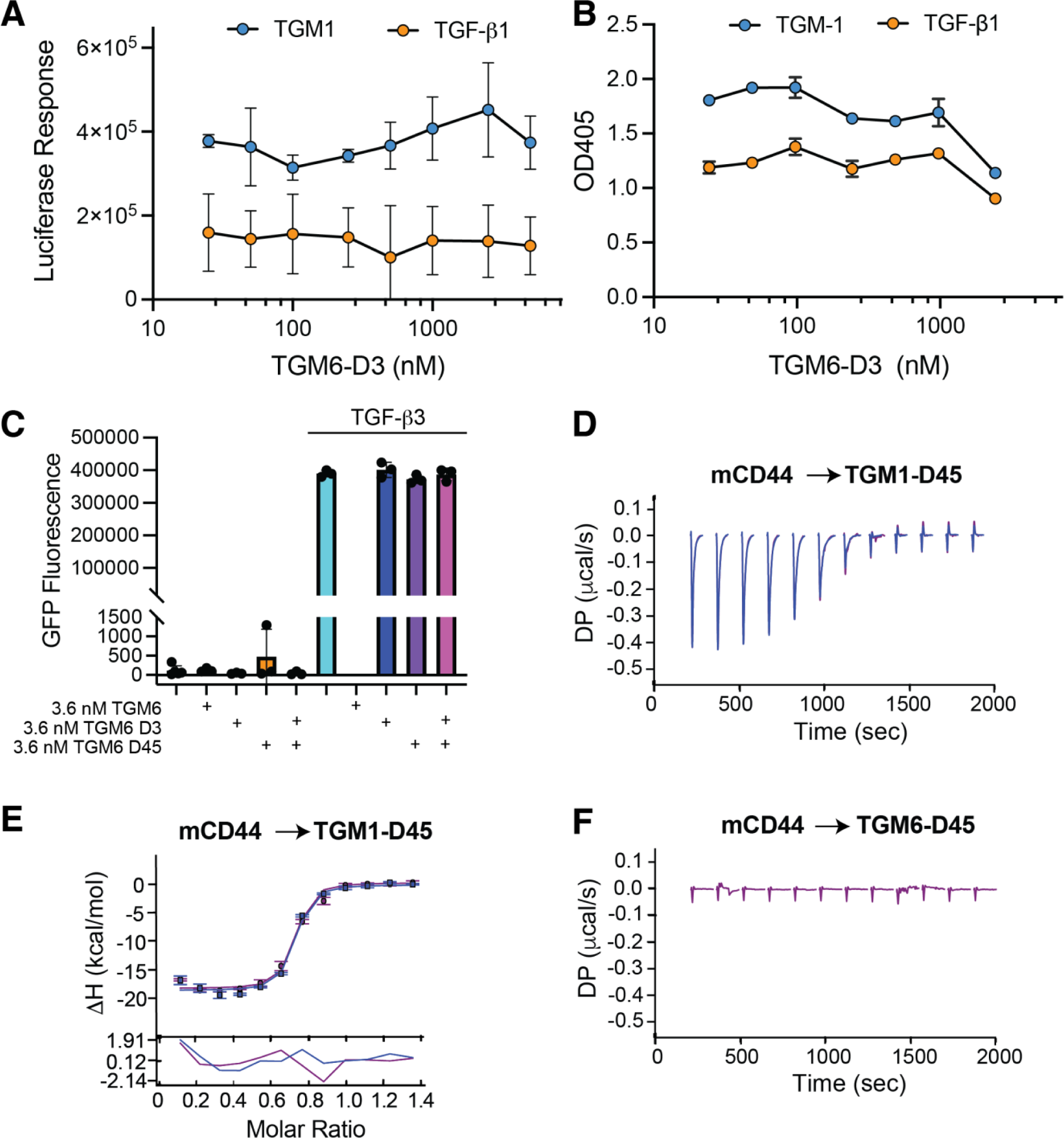
TGM6 requires attachment of D3 to D45 to inhibit and does not bind the TGM1 co-receptor CD44. **A-B.** Inhibition of TGF-β reporter stimulated by TGF-β1 or TGM1 in NIH-3T3 (**A**) or MFB-F11 (**B**) fibroblasts by increasing concentrations of TGM6-D3. Data could not be reliably fit to a dose-dependent inhibition model. **C.** Inhibition of TGF-β GFP reporter in NIH-3T3 fibroblasts by TGM6, TGM6-D3, TGM6-D45, or TGM6-D3 plus TGM6-D45. **D-E**. ITC thermogram (**D**) and fitted isotherm (top) and residuals (bottom) (**E**) for titration of mCD44 into TGM1-D45. **F**. ITC thermograms for titration of mCD44 into TGM6-D45.

The inability to TGM6 to inhibit signaling in splenic T cells, together with the finding that TGM1 constructs lacking the CD44 co-receptor binding domains attenuated the ability of TGM1 to convert naïve T cells to Foxp3^+^ Tregs [31], suggested that domains 4 and 5 of TGM6 may not bind CD44. To investigate this, we used ITC to measure the binding affinity of TGM6-D45, and as a control TGM1-D45, for mouse CD44 (mCD44) (**Fig. 5D-F**). The titration of mCD44 into TGM6-D3 led to an insignificant response that did not change over the course of the titration, while titration of mCD44 into TGM1-D3 led to a robust exothermic response, which upon integration, yielded a K_D_ in close accord with that reported earlier [31] (**Table S4**). TGM6’s dependence on domains 4 and 5 for inhibition, and the inability of TGM6-D45 to bind CD44, are therefore likely responsible for its inability to inhibit TGF-β1 and TGM1 signaling in T cells.

### TGM6-D3 mimics TGF-β binding to TβRII

To identify the underlying molecular basis by which TGM6-D3 binds and recognizes TβRII, we isolated the TGM6-D3:TβRII complex using size exclusion chromatography, screened for diffracting crystals, and determined the structure to a resolution of 1.40 Å (**Fig. 6A, Table S5**). We found one TGM6-D3:TβRII complex in the crystallographic asymmetric unit and interpretable density for residues 46-153 of TβRII and residues 16-65, 71-82, and 86-102 of TGM6-D3. Overlay of the TGM6-D3-bound TβRII structure with the previous crystal structure of unbound TβRII (PDB 1M9Z) [36] revealed only minor differences, with a backbone RMSD of 0.49 Å overall (**Fig. S7A**). Overlay of the TβRII-bound TGM6-D3 structure with the lowest energy unbound TGM1-D3 structure determined by NMR (PDB 7SXB) [28] revealed close similarity, with a backbone RMSD of 1.25 Å overall (**Fig. S7B**). Residues missing in the density for TGM6-D3, Lys^66^–Ala^70^ and Arg^83^–Thr^85^, correspond to the tips of the loops connecting β-strands 2-3 and 3–4, which were previously shown to be flexible in unbound TGM1-D3 [28].

**Figure 6.**
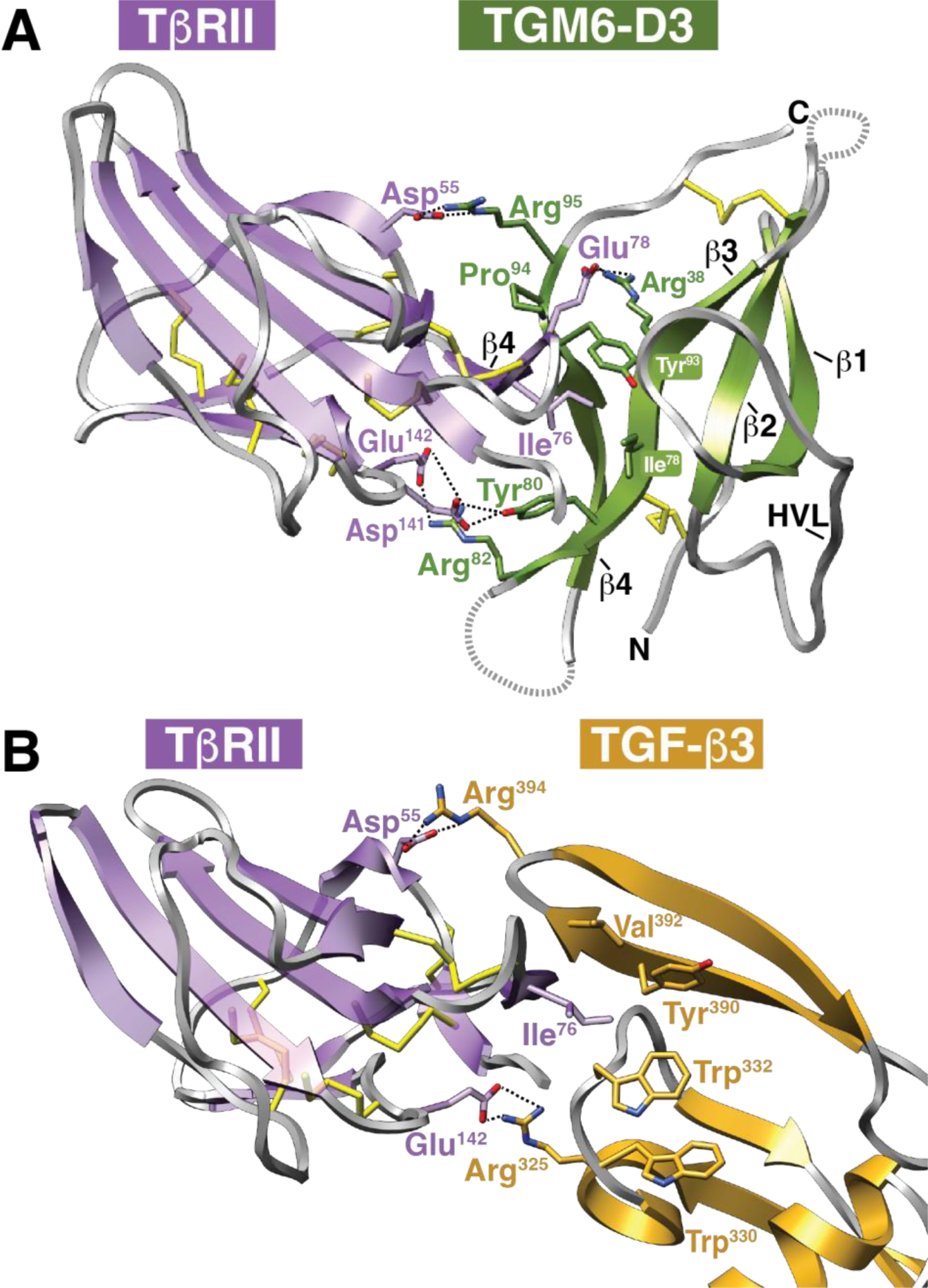
Structure of the TGM6-D3:TβRII complex and mimicry of mammalian TGF-β. **A.** Overall structure of the TGM6-D3:TβRII complex determined by X-ray crystallography at a resolution of 1.4 Å. TGM6-D3 and TβRII are shaded olive green and lavender, respectively. Loops that are not modeled due to weak density are indicated by dashed lines. Sidechains of key interfacial residues are shown, as are the intramolecular disulfide bonds. **B.** Structure of the TGF-β3:TβRII complex at a resolution of 2.15 Å (PDB 1KTZ). TGF-β1 and TβRII are shaded burnt orange and lavender, respectively. Sidechains of key interfacial residues are shown, as are the intramolecular disulfide bonds.

There is a single large interface between TGM6-D3 and TβRII in the lattice (interface area of 661 Å^2^) that has an overall manner of binding in accord with that anticipated based on the previous studies of TGM1-D3 [28] (**Fig. 6A**). TGM6-D3 engages TβRII primarily through the edge β-strand, β4, and does so through the interface near the C-terminus with the residues anticipated, Ile^78^ from β-strand 3, Tyr^93^ and Arg^95^ from β-strand 4, and Arg^38^ from the tip of the hypervariable loop (HVL) (**Fig. 1C-D**). The interface between TGM6-D3 and TβRII is a remarkable mimic of the interface between TGF-β1/-β3 and TβRII [37, 38], with a central hydrophobic region flanked by hydrogen-bonded ion-pairs at the periphery (**Fig. 6**). On one side of the interface, TβRII Asp^55^ and Glu^78^ form hydrogen-bonded ion-pairs with TGM6-D3 Arg^95^ and Arg^38^, respectively (**Fig. 6A, 7A**), the first of which closely mimics the interaction between TβRII Asp^55^ and TGF-β1/-β3Arg^394^ (**Fig. 6B, 7B**). In the central hydrophobic region, TβRII Ile^76^ inserts into a hydrophobic pocket formed by TGM6-D3 residues Ile^78^, Tyr^80^, and Tyr^93^ (**Fig. 6A, 7C**), mimicking the interaction between TβRII Ile^76^ and the hydrophobic pocket between the fingers of TGF-β formed by Trp^332^, Tyr^390^, and Val^392^ (**Fig. 6B, 7D**). On the side of the interface opposite the TGM6-D3 Arg^95^:TβRII Asp^55^ interaction, TβRII Asp^141^ forms a hydrogen bond with the phenolic hydroxyl of TGM6-D3 Tyr^80^ and TβRII Glu^142^ forms a hydrogen-bonded ion-pair with TGM6-D3 Arg^82^ (**Fig. 6A, 7E**), mimicking the hydrogen-bonded ion-pair between TβRII Glu^142^ and TGF-β1/-β3 Arg^325^ (**Fig. 6B, 7F**).

### The interactions that enable TGM6-D3 and TGF-β1/-β3 to bind TβRII are similar

Through mutagenesis and binding studies, the central hydrophobic interaction and two flanking hydrogen-bonded ion-pairs in the interface between TGF-β1/-β3 and TβRII were each shown to contribute significant to binding [39, 40]. Though the overall architecture of the TGM6-D3:TβRII interface is similar to the TGF-β1/-β3:TβRII interface, it is about 50% larger (661 Å^2^ vs. 479 Å^2^ for the TGF-β1/-β3:TβRII interface) and it has a greater number of hydrogen bond and ionic interactions (**Fig. 6**). Thus, to assess the contributions of these interactions in the TGM6-D3:TβRII complex, we substituted single residues in both TGM6-D3 and TβRII and used ITC to characterize the binding (**Fig. S8, Table S6**).

To analyze the TGM6-D3 Arg^95^:TβRII Asp^55^ doubly hydrogen-bonded ion-pair (**Fig. 7A**), we substituted TβRII Asp^55^ or TGM6-D3 Arg^95^ with alanine and found that these diminished the binding affinity by 6.4- or 11.6-fold (**Table S6**). There is an additional nearby ion-pair between TβRII Glu^78^ positioned in the loop following β-strand 4 and TGM6-D3 Arg^38^ on the tip of the HVL and substitution of TGM6-D3 Arg^38^ with alanine diminished the binding affinity by 22-fold. If it is assumed the binding energy in this region of the TGM6-D3:TβRII complex is a result of both the interactions described above, then the theoretical perturbation of simultaneously eliminating both interactions would be 140-255-fold. In the TGF-β1/-β3:TβRII complex substitution of the residues that form the homologous doubly hydrogen-bonded ion-pair, TβRII Asp^55^ and TGF-β1/-β3 Arg^394^, led to a 30- to 80-fold decrease in binding affinity (**Fig. 7B, Table S6**). Thus, the energetic contribution of the two combined interactions in the TGM6-D3:TβRII complex are comparable or exceed the contribution of the single interaction in the TGF-β1/-β3:TβRII complex.

**Figure 7.**
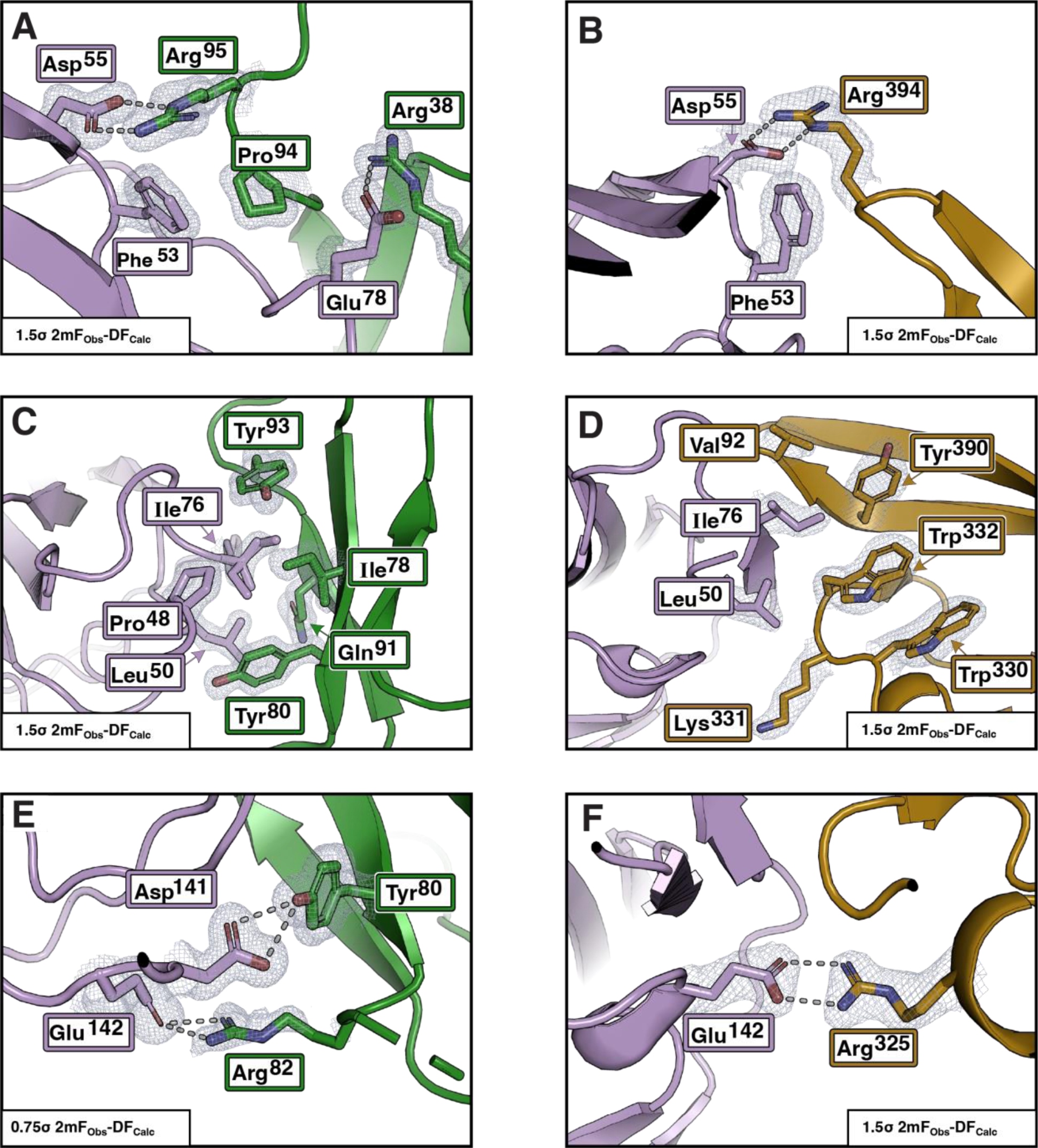
Interface contacts of TGM6-D3 with TβRII and mimicry of mammalian TGF-β. **A-B.** Ionic interaction with TβRII Asp^55^ with Arg^95^ of TGM6-D3 (**A**) and Arg^94^ of TGF-β3 (**B**). TβRII Glu^78^ also has an ionic interaction with TGM6-D3 Arg^38^, but it does not interact with TGF-β. **C-D.** Interaction of TβRII Leu^50^ and Ile^76^ with the hydrophobic pocket on TGM6-D3 formed by Tyr^93^, Ile^78^, and Tyr^80^ (**C**) and TGF-β3 formed by Tyr^90^, Val^92^, and Trp^30^ (**D**). **E-F**. Ionic interaction with TβRII Glu^142^ and Asp^141^ with Tyr^80^ and Arg^82^ of TGM6-D3, respectively (**E**) and TβRII Glu^142^ with Arg^25^ of TGF-β3 (**F**). In all panels, TβRII is shaded lavender and sidechains of key interaction residues are shown. In panels **A**, **C**, and **E** TGM6 is shaded green and in panels **B**, **D**, and **F** TGF-β3 is shaded burnt orange.

The central hydrophobic interaction between TβRII and TGM6-D3 was probed by substituting both TβRII Ile^76^ and TGM6-D3 Ile^78^, Tyr^80^, and Tyr^93^ with alanine (**Fig. 7C**). These substitutions led to large perturbations, ranging from 21-fold for the TβRII I76A substitution, to 15.4-, 134-, and 108-fold for the TGM6-D3 I78A, Y80A, and Y93A substitutions, respectively (**Table S6**). In the TGF-β3:TβRII complex, substitution of Ile^76^ with alanine led to a 12.5-fold reduction in affinity, while the effects of substituting of Trp^30^, Tyr^90^, and Ile^92^ between the fingers of the TGF-β1/-β3 was not reported due to the propensity of substitutions in this region of the protein to lead to misfolding [41] (**Fig. 7D, Table S6)**. The methyl and amide regions of the 1D ^1^H NMR spectra of the TGM6-D3 I78A, Y80A, and Y93A variants have dispersed patterns similar to wild type, indicating that the effects of the substitutions on binding is not due to perturbation of the folding (**Fig. S9**). The central hydrophobic interaction clearly has an essential role for binding in the TGM6-D3:TβRII complex, similar to that in the TGF-β1/- β3:TβRII complex, however it is not possible to directly compare the energetic contribution of these to binding due to the lack of TGF-β mutants and the multiple residues involved.

The functional significance of the TGM6-D3 Arg^82^:TβRII Glu^142^ hydrogen-bonded ion-pair was uncertain as there was defined, but weaker electron density for the Arg^82^ guanidinium group, but little density for the sidechain Cβ and Cγ atoms and the backbone (**Fig. 7E**). The modeling of the Arg^82^ guanidinium group is supported by omit maps in which Arg^82^ and the subsequent three residues in the β3-β4 loop are absent (**Fig. S10**); further, substitution of TGM6-D3 Arg^82^, or its partner residue, TβRII Glu^142^, with alanine is shown to diminish the binding affinity by 11- to 12-fold (**Table S6**). The adjacent hydrogen-bonding interaction between the phenolic hydroxyl of TGM6-D3 Tyr^80^ and the sidechain carboxylate of Asp^141^ also appears to be functionally significant as substitution of Asp^141^ to alanine or TGM6-D3 Tyr^80^ with phenylalanine reduced the affinity by 2.7- and 3.5-fold. Thus, similar to TGM6-D3 Arg^95^:TβRII Asp^55^ and TGM6-D3 Arg^38^:TβRII Glu^78^ ion-pairs on the opposite side of the interface, both interactions appear contribute and the theoretical perturbation of simultaneously eliminating both interactions is expected to be about 35-fold. In the TGF-β1/-β3:TβRII complex substitution of the residues that form the homologous doubly hydrogen-bonded ion-pair, TβRII Glu^142^ and TGF-β1/-β3 Arg^325^, led to a 12- to 30-fold decrease in the binding affinity (**Fig. 7F, Table S6)**. Thus, the combination of the hydrogen-bond and ion-pair interactions in TGM6-D3:TβRII complex and hydrogen-bonded ion-pair in the TGF-β1/-β3:TβRII complex contribute comparably to the overall binding energy.

### TGM6 function is highly dependent on high affinity TβRII binding, while TGM1 function is not

TGM1 is dependent upon both TβRI and TβRII for signaling, but it binds TβRII moderately (K_D_ 1.2 -1.5 µM vs. 0.35 µM for TGM6) and single amino acid substitutions, such as Y253A that increase the K_D_ for TβRII by more than 40-fold, only modestly attenuate signaling [31]. In contrast, the Y253A substitution nearly abrogates signaling, even with very high concentrations of ligand, when the co-receptor binding domains, D4-D5, are absent [28]. Thus, when there is trivalent engagement of receptors, TGM1 signaling activity is retained even if TβRII binding affinity is significantly compromised, but when there is bivalent engagement of receptors, the same perturbation of TβRII binding almost eliminates signaling activity.

This suggested that identification of TGM1-D3 residues responsible for its lower affinity for TβRII and swapping these into TGM6-D3 would severely impair its inhibitory potential, owing to its bivalent engagement of receptors, TβRII through D3 and a co-receptor through D4-D5. The substitution of the corresponding TGM6-D3 residues into TGM1-D3, by contrast, was expected to only modestly enhance signaling owing to its trivalent engagement of receptors.

To investigate this, we compared the amino acid sequences of TGM1-D3 and TGM6-D3, in which homologous residues differ in notation by the 160-amino acid length of D1-D2 present only in TGM1. We noted two differences that might be responsible for the reduced affinity of TGM1 for TβRII (**Fig. 1D**). The first is substitution of the Lys^254^-Asn^255^ (KN) dipeptide in TGM1 in place of the Pro^94^-Arg^95^ (PR) dipeptide in TGM6. The second is substitution of the Lys^241^-Ser^242^-Gly^243^-Thr^244^ (KSGT) tetrapeptide in TGM1 in place of the Gln^81^-Arg^82^-Arg^83^-Gly^84^ (QRRG) tetrapeptide in TGM6. The Lys^254^ or Lys ^241^ of TGM1 KN or KSGT peptides may not functionally replicate the hydrogen-bonded ion-pairs of TGM6 Arg^95^ or Arg^82^ with TβRII Asp^55^ or Glu^142^, respectively, leading to weaker binding and impairment of function (**Fig. 7A, 7E**).

The first of these was tested by substituting the PR dipeptide of TGM6-D3 with the KN dipeptide from TGM1, and vice versa, and measuring the binding affinity of the chimeric proteins, TGM6-D3 KN and TGM1-D3 PR, for TβRII using ITC (**Fig. S8**). Unexpectedly, the affinities of TGM1-D3 PR and TGM6-D3 KN for TβRII were either slightly weaker or indistinguishable from their parental wild type proteins, indicating that the lysine of the KN dipeptide of TGM1 can functionally replicate the interaction that TGM6-D3 Arg^95^ has with TβRII Asp^55^ (**Table S6**). The second was tested by exchanging the KSGT tetrapeptide in TGM1 with the QRRG tetrapeptide in TGM6 and vice versa, again measuring the affinity of these chimeric proteins for TβRII using ITC (**Fig. 8A-B**). These showed that binding of TGM6-D3 KSGT to TβRII was impaired by 12.5-fold compared to wild type TGM6-D3, while binding of TGM1-D3 QRRG to TβRII was enhanced by 8.5-fold compared to wild type TGM1-D3, indicating that these residue differences indeed underlie the differential affinity of TGM1-D3 and TGM6-D3 for TβRII. To ascertain whether replacement of TGM6-D3 Arg^82^ with the corresponding residue of TGM1-D3, Ser^242^, and vice versa were responsible for the loss and gain of affinity, we generated the TGM6-D3 R82S and TGM1-D3 S242R single amino acid variants and measured their affinity for TβRII using ITC (**Fig. S8, Table S6**). The loss and gain of binding affinity compared to wild type were more moderate compared to the tetrapeptide swaps, 7.4-fold loss for TGM6-D3 R82S and 4.0-fold gain of for TGM1-D3 S242R, but they were also more aligned with the overall affinity difference between TGM1 and TGM6 for TβRII, indicating that these alone are responsible for the affinity difference.

**Figure 8.**
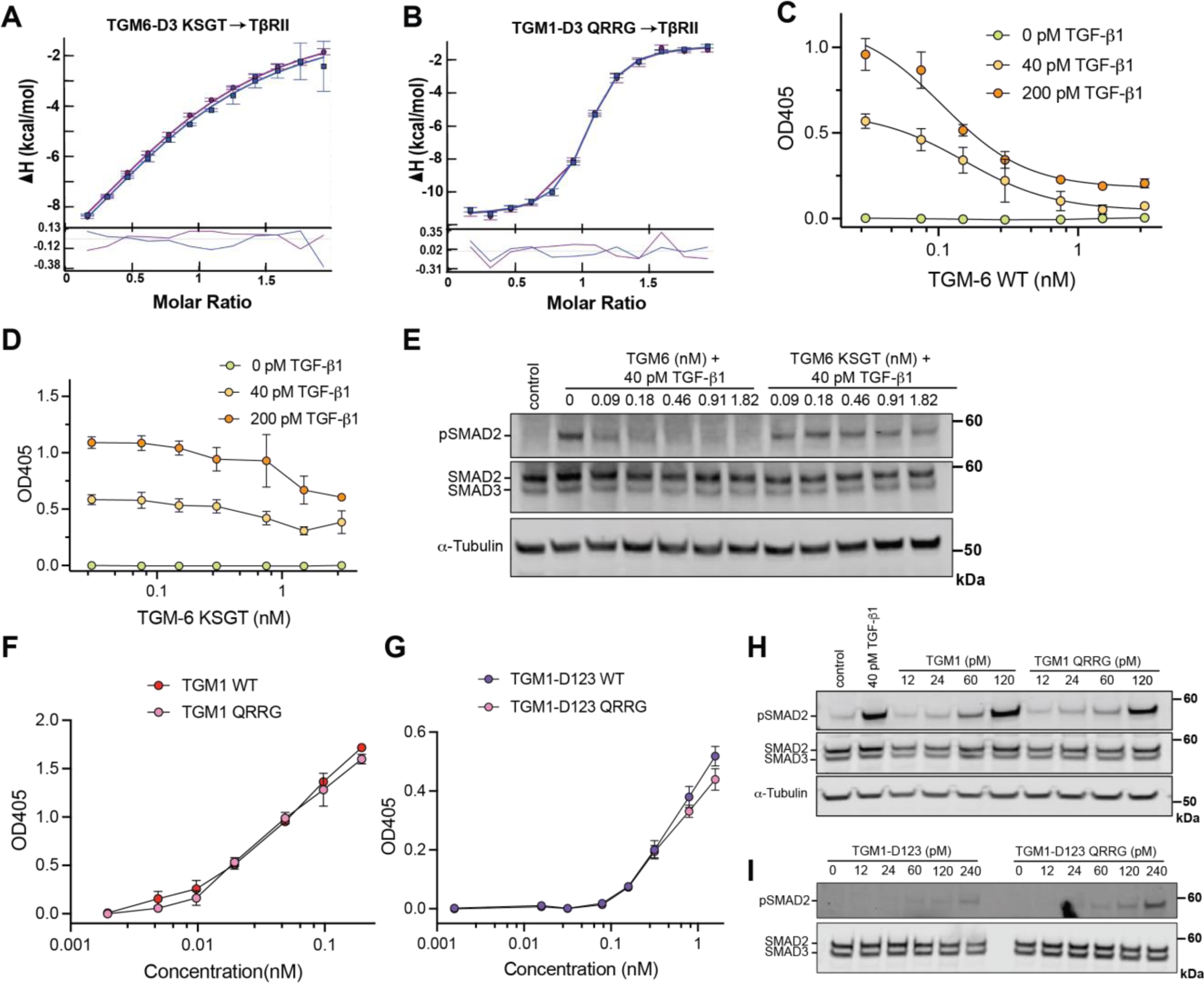
TGM6 residues responsible for its high affinity for TβRII. **A-B.** ITC fitted binding isotherms (top) and residuals (bottom) for titration of TGM6-D3 KSGT (**B**) or TGM1-D3 QRRG (**C**) into TβRII. **C-E.** Inhibition of TGF-β1 signaling in MFB-F11 fibroblasts as detected by the conversion of p-nitrophenylphosphate by secreted alkaline phosphatase (**D-E**) or by pSMAD2 Western blotting (**E**) by either TGM6 (**C, E**) or TGM6 KSGT (**D, E**). **F-I**. Activation of TGF-β1 signaling in MFB-F11 fibroblasts as detected by the conversion of p-nitrophenylphosphate by secreted alkaline phosphatase (**F-G**) or by pSMAD2 Western blotting (**H-I**) by either TGM1 or TGM1 QRRG (**F, H**) or by TGM1-D123 or TGM1-D123 QRRG (**G, I**).

To determine how the loss or gain of TβRII binding affinity affected function, we began by comparing the inhibitory activity of TGM6 or TGM6 KSGT in MFB-F11 fibroblasts as assessed by either its secreted alkaline phosphatase reporter activity or induction of pSMAD2/3. In both assays TGM6 KSGT failed to inhibit induction of signaling by either TGF-β1 or TGM1, except at the highest concentration where it only partially inhibited, while TGM6 WT inhibited at all concentrations tested (**Fig. 8C-E, Fig. S11**). We then compared the induction of signaling in MFB-F11 fibroblasts by TGM1 or TGM1-D123, either as the wild type protein or the QRRG variant. In the reporter assays, we observed no apparent difference between the wild type protein or QRRG variant, regardless of whether the full-length or truncated protein was assayed (**Fig. 8F-G**), while in the pSMAD2/3 assays, we observed no apparent differences between the wild type protein or QRRG variant in the context of full-length TGM1 (**Fig. 8H**), but a small but consistent increase in potency at the three highest concentrations tested in the context of the truncated protein (**Fig. 8I**). Thus, consistent with expectations, swapping the residues responsible for the lower affinity of TGM1 for TβRII into TGM6 dramatically impaired its inhibitory activity, whereas swapping the residues responsible for the greater affinity of TGM6 for TβRII into TGM1 only modestly enhanced its signaling activity.

TGM2 and TGM3, like TGM1, retain the key residues identified in TGM6 required for high affinity TβRII binding, but lack a basic residue homologous to TGM6 Arg^82^, likely accounting for the comparable signaling activity of TGM1-3 in MFB-F11 fibroblasts (**Fig. S12**) [29]. TGM4 which binds TβRII more weakly than TGM1 [32] and does not signal in MFB-F11 fibroblasts [29], also lacks a basic residue homologous to TGM6 Arg^82^, but also absent is the Pro^94^-Arg^95^ dipeptide of TGM6 or the Lys^254^-Asn^255^ dipeptide of TGM1. TGM7, which has not been found to have any TGF-β signaling activity in MFB-F11 fibroblasts [29], or TGM8, which has not functionally characterized, retain an arginine homologous to TGM6 Arg^82^, but have either a His-Asn or Leu-Asn dipeptide in place of the KN dipeptide of TGM1 or the PR dipeptide of TGM6. TGM5, which lacks domain 4, TGM9 which lacks domains 1 and 2, and TGM10 which lacks domains 3 and 4 have not been functionally characterized. TGM9 retains all key residues identified in TGM6 that contribute to high affinity TβRII binding. Thus, TGM9 might also inhibit TGF-β signaling, which we are currently investigating.

## DISCUSSION

The persistence of the helminth *H. polygyrus* in its mammalian host is dependent upon the upregulation of Tregs [11], and as shown this activity stems from TGM1, and likely other five domain TGM homologs such as TGM2, 3, and 4, that bind the host TGF-β receptors, TβRI and TβRII, and activate the TGF-β pathway [6, 29]. Though lacking any sequence or structural homology to the TGF-βs, TGMs recognize the same regions of TβRI and TβRII as TGF-β, suggesting that they are not only functional mimics of mammalian TGF-β, but also structural mimics [28].

TGMs have also adapted in other surprising ways to successfully evade host immunity. TGM1, for example, was shown to bind the co-receptor CD44 through domains 4 and 5 [31]. This increases the signaling potency and biases TGM1 activity to cells which have abundant CD44, such as T lymphocytes and myeloid cells [31, 32]. TGM4, which shares the same five-domain structure as TGMs 1, 2, and 3, was recently shown to bind TβRI 10-fold more strongly than TGM1, but TβRII 100-fold less avidly than TGM1 [32]. This combination of receptor binding affinities, together with binding of CD44 and several other co-receptors, including CD206, Itga4, and neuropilin-1, biases TGM4 activity toward different subclasses of myeloid cells that express these co-receptors.

The studies of TGM6 reported here further contribute to our understanding of the adaptations of the TGM family of proteins. The structure of TGM6-D3 bound to TβRII reveals the remarkable molecular mimicry that enables domain 3 to bind TβRII in a manner that closely resembles TGF-β1/- β3 [37, 38]. In the structure of the TGM6-D3:TβRII complex, we find that not only does the parasite protein engage TβRII in the same overall manner as TGF-β1/-β3 through a central hydrophobic interaction and flanking hydrogen-bonded ion-pairs, but the presentation of key interacting residues, such as Arg^95^ and Arg^82^, which form hydrogen-bonded ion-pairs with TβRII Asp^55^ and Glu^142^, are also remarkably similar to those of TGF-β1/-β3. Though the hydrogen-bonded ion-pairs that TGM6-D3 Arg^95^ and Arg ^82^ have with Asp^55^ and Glu^142^ of TβRII do not contribute as much energetically as the corresponding interactions in the TGF-β1/-β3 complex, this is compensated by additional hydrogen-bonds and hydrogen-bonded ion-pairs, including TGM6-D3 Arg^38^:TβRII Glu^78^ and TGM6-D3 Tyr^80^:TβRII Asp^141^, that are lacking in the TGF-β1/-β3:TβRII complex. The additional interactions in the TGM6-D3:TβRII complex arise in part from adaptations of the TGM-D3 CCP fold, such as the structurally ordered HVL [28], that provide a greater number of opportunities to position residues that can productively interact with residues on TβRII.

This study has also shown that among the TGMs secreted by the adult parasite, the three-domain TGM6 retains the capacity to bind TβRII through domain 3, but it does not bind TβRI or activin and BMP type I and type II receptors, and thus functions as a potent inhibitor of TGF-β and TGM1 signaling by competitively binding TβRII. The paradox of why the adult parasite might have evolved to co-express inhibitory TGM6 alongside signaling-activating TGMs likely stems from the different cell populations they target, with TGM6 potently inhibiting TGF-β and TGM1 signaling in fibroblasts, but not splenic T cells, the cell type targeted by TGM1 [31]. The targeting of TGM6 to fibroblasts is likely mediated by binding of domains 4 and 5 to a co-receptor that is present on fibroblasts, but not T cells. This is suggested by the finding that a) domain 3 alone is not inhibitory, b) the inhibitory potency of the full-length protein is 1300-fold lower than the binding affinity of domain 3 for TβRII, and c) domains 4 and 5 of TGM6 do not bind CD44.

The targeting of TGM6 to fibroblasts might reflect a strategy the parasite has adopted to reduce fibrotic activity as it transitions through its life cycle, which involves invasion of newly arriving larvae through the intestinal epithelium to encyst in the muscle. Upon maturing in the intestinal submucosa, the adult parasites burrow back through the intestinal wall to the lumen where they mate and produce eggs that are released to the environment in the feces. In light of the considerable tissue damage that would occur in this process, and the well-established role of TGF-β and TGM1 in contributing to wound repair by stimulating deposition of type I collagen (16, 33, 54), but also driving tissue fibrosis if signaling is dysregulated (16), it is possible that TGM6 modulates TGF-β signaling in fibroblasts, which would in turn reduce the degree of collagen deposition and fibrosis.

The TGF-βs, in addition to stimulating pro-tolerogenic signaling that is required for immune homeostasis, are highly pleiotropic and induce signaling that is important for homeostatic control of many other essential processes [42]. The targeting of the TGF-β pathway by the helminth *H. polygyrus* therefore has potentially adverse consequences for the host. To overcome potentially deleterious off-target effects, and thus limit damage to the host, the parasite has adapted the multidomain CCP scaffold proteins, not only to bind TβRI and TβRII to activate the TGF-β pathway, but also to bind co-receptors that precisely target them to relevant immune cell subsets [6, 31, 32].

A further evolutionary adaptation has been calibrating the affinities of the relevant receptor binding domains to confer target specificity, achieved by weakening binding to TβRII and by enhancing binding to the relevant co-receptors present on different immune cell subsets [28, 31, 32]. In TGM agonists with trivalent engagement of receptors, TβRI, TβRII, and a co-receptor, it is evidently possible to severely attenuate TβRII binding, while still effectively assembling TβRI:TβRII complex for signaling [32]. In antagonists, such as TGM6, with bivalent engagement of receptors, TβRII and a co-receptor, it is not possible to weaken TβRII binding to the same extent as the agonists. In spite of this, it appears the parasite has tuned the binding affinity of TGM6-D3 for TβRII to enable effective antagonism only when also bound to a co-receptor through domains 4 and 5, allowing it to discriminate between target populations according to their co-receptor profile.

In summary, our understanding of the *H. polygyrus* TGM family of proteins is expanded by showing that TGM6 is a potent inhibitor of TGF-β- and TGM1-induced TGF-β signaling in fibroblasts, and that it does so without interfering with essential immune-suppressive signaling of TGM agonists in T cells. The structure of the TGM6-D3:TβRII binary complex shows remarkable mimicry of the TGF- β:TβRII interactions and the accompanying functional studies not only pinpoint the residues responsible for the greater affinity of TGM6 domain 3 for TβRII compared to TGM1 domain 3, but also highlight how the TGMs tune their affinity for TβRII to direct them only to immune cells and fibroblasts in which their cognate co-receptor(s) is (are) expressed.

## MATERIALS AND METHODS

### Expression and purification of TGM proteins

The amino acid sequences of the TGM proteins used in this study are presented in **Table S7**. The plasmids used to produce TGM1, TGM1-D123, and TGM6 in mammalian cells were previously described [6, 29]. The plasmids used to produce TGM1-D3, TGM6-D3, and TGM6-D45 in bacteria were generated by inserting the corresponding coding sequences downstream of the thrombin cleavage site in a modified form of pET32a (EMD-Millipore) with a His_10_ tag instead of His_6_.

TGM1, TGM1-D123, and TGM6 were expressed in expi293 cells (Invitrogen) and purified by binding the protein in the conditioned medium onto a nickel-loaded chelating Sepharose column (Cytiva). The bound protein was eluted using a 0 – 0.5 M imidazole gradient. The eluted protein peak was concentrated, deglycosylated with a 14 h incubation with PNGAse F at 30°C, and further purified by size exclusion chromatography (HiLoad 26/60 Superdex 75 column, Cytiva).

TGM1-D3 was overexpressed in *E. coli* at 37°C in the form of insoluble inclusion bodies, refolded, and purified as described [43]. TGM6-D3 and TGM6-D45 were produced and purified similarly, with two exceptions: 1) the lysis supernatant for TGM6-D3 was retained and combined with the urea-solubilized inclusion bodies prior to purification by nickel metal affinity chromatography and 2) rather than final purification on a Source Q column, both were purified on a Source S column (Cytiva). TGM6-D3 was bound in 25 mM sodium acetate, 2 M urea, 10 μM leupeptin hemisulfate, 10 μM pepstatin, 100 mg L^-1^ benzamidine, pH 4.8 and eluted with a 0 – 0.35 M NaCl gradient, while TGM6-D45 was bound in 25 mM Tris, 2 M urea, 10 μM leupeptin hemisulfate, 10 μM pepstatin, 100 mg L^-1^ benzamidine pH 7.5, and eluted with a 0 – 0.2 M NaCl gradient.

### Expression and purification of type I receptors

The amino acid sequences of the type I receptors used in this study are presented in **Table S8**. The plasmids used to produce ALK1, ALK3, and ALK5 (also known as TβRI) in bacteria were previously described [43, 44]. The plasmids used to produce ALK2 and ALK4 in bacteria were constructed by inserting the corresponding coding sequence downstream of the thrombin cleavage site in either pET32a or pET15b (EMD-Millipore).

ALK1, ALK3, and ALK5 were expressed in *E. coli* grown on M9 minimal medium with ^15^N-labeling, refolded from urea-solubilized inclusion bodies, and purified as described [43, 44]. ALK4 was expressed, refolded, and purified similarly to ALK1, with the exception that the refolded protein was purified in two steps, first by loading onto a Source Q column in 25 mM Tris, 2M urea, 10 μM leupeptin hemisulfate, 10 μM pepstatin, and 100 mg L^-1^ benzamidine pH 8.0 and eluting with a 0 – 0.35 M NaCl gradient, and second by loading onto a C18 semi-preparative reverse phase column (Jupiter 5 μm C18 300 Å; Phenomenex) and eluting with a 5 - 70% acetonitrile gradient.

ALK2 was expressed on minimal medium with ^15^N labeling at 37°C until the A_600_ reached 0.4, followed by transfer to 14°C and induction of expression with IPTG when the A_600_ reached 0.6. ^15^N-ALK2 protein was purified from the lysis supernatant by loading onto a nickel-loaded chelating Sepharose column (Cytiva) and eluting with a 0 - 0.5 M imidazole gradient. Fractions containing ^15^N-ALK2 were pooled, thrombin treated, dialyzed against 25 mM CHES, pH 9.0, and purified in two steps, first by loading onto a Source Q column (Cytiva) and eluting with a 0 – 0.35 M NaCl gradient, and second by loading onto a C18 semi-preparative reverse phase column (Jupiter 5 μm C18 300 Å, Phenomenex) and eluting with a 5 - 70% acetonitrile gradient.

### Expression and purification of type II receptors

The amino acid sequences of the type II receptors used in this study are presented in **Table S9**. The plasmid used to produce TβRII in bacteria was previously described [45]. The plasmids used to produce ActRII and ActRIIB in bacteria were constructed by inserting the corresponding coding sequence downstream of the N-terminal His_6_ tag and thrombin cleavage site in pET15b (EMD-Millipore). The plasmid used to produce BMPRII extracellular domain in mammalian cells was constructed by inserting the coding sequence for this downstream of the rat serum albumin signal peptide, a His_6_ tag, and a thrombin cleavage site in a modified form of pcDNA3.1+ (Invitrogen).

TβRII was overexpressed in *E. coli* at 37°C in the form of insoluble inclusion bodies, refolded, and purified as described previously [45]. ActRII and ActRIIb were expressed in the form of insoluble inclusion bodies in *E. coli* BL21(DE3) cells (EMD-Millipore) and were refolded and purified similarly to ALK1, with the exception that the refolded protein was purified by loading onto a Source Q column in 25 mM sodium phosphate, 10 μM leupeptin hemisulfate, 10 μM pepstatin, and 100 mg L^-1^ benzamidine pH 6.6 and eluting with a 0 – 0.35 M NaCl gradient. BMPRII was expressed in expi293 cells (Invitrogen) and purified from the conditioned medium using nickel metal affinity chromatography and SEC in the manner described above for TGM6.

### Expression and purification of mmTGF-β2-7M2R and mCD44

The amino acid sequences of the miniTGF-β, mmTGF-β2-7M2R, and mouse CD44 (mCD44) used in this study are presented in **Table S10**. mm-TGF-β2-7M2R was overexpressed in *E. coli* BL21(DE3) cells at 37°C in the form of inclusion bodies, refolded, and purified as previously described [35]. mCD44 was produced in expi293 suspension cultured mammalian cells and purified as previously described [31].

### Point mutants and validation of recombinant proteins

Single amino acid mutants of TβRII, TGM6-D3, or TGM1-D3 were generated by site-directed mutagenesis [46]. Multiple amino mutants of TGM1 and TGM6 were generated by gene synthesis (Twist Biosciences). Coding sequences of all wild type and mutant proteins were verified by DNA sequencing over the length of their coding sequences. Masses of all recombinant proteins produced in *E. coli*, including all point mutants, were confirmed by liquid chromatography electrospray ionization time-of-flight mass spectroscopy (Micro TOF, Bruker).

### NMR data collection

Samples of ^15^N TGM6-D3 and its complex with TβRII were prepared at a concentration of 150 μM in 25 mM Na_2_HPO_4_, 10 μM leupeptin hemisulfate, 10 μM pepstatin, 100 mg L^-1^ benzamidine, 0.05% (w/v) NaN_3_, 5% ^2^H_2_O, pH 5.5. Samples of ^15^N ALK1, ^15^N ALK2, ^15^N ALK3, ^15^N ALK4, and ^15^N ALK5 and their corresponding samples containing 1.125 molar equivalents of TGM6-D45 or the TGM6:TβRII binary complex were prepared at a concentration of 100 μM ^15^N-labelled receptor in 25 mM Na_2_HPO_4_, 10 μM leupeptin hemisulfate, 10 μM pepstatin, 100 mg L^-1^ benzamidine, 0.05% (w/v) NaN_3_, 5% ^2^H_2_O, pH 6.0.

All NMR samples were transferred to 5 mm susceptibility-matched microtubes for data collection (Sigma-Aldrich). NMR data were collected at 303.15 K using 600, 700, or 800 MHz spectrometers equipped with 5 mm ^1^H (^13^C,^15^N) z-gradient “TCI” cryogenically cooled probes (Bruker Biospin). 2D ^1^H-^15^N HSQC spectra were recorded with sensitivity enhancement [47], water flip-back pulses [48], and WATERGATE water suppression pulses [49]. NMR data were processed using NMRPipe [50] and analyzed using NMRFAM-SPARKY [51].

### ITC measurements

ITC data was generated using a Microcal PEAQ-ITC instrument (Malvern Instruments, Westborough, MA). All experiments were performed in ITC buffer (25 mM HEPES, 150 mM NaCl, 0.05% NaN_3_, pH 7.4). The proteins in the syringe and sample cell and their concentrations are provided in the respective data tables. Prior to each experiment, all proteins were dialyzed three times against ITC buffer and were concentrated or diluted as necessary before being loaded into the sample cell or syringe. For each experiment, nineteen 2.0 μL injections were performed with an injection duration of 4 sec, a spacing of 150 sec, and a reference power of 10. Integration and data fitting were performed using Nitpic [52] and Sedphat [53, 54]. No more than two outlier data points were removed from any one ITC data set for analysis. The TGM6:TβRII binding experiment was globally fit to a simple binding model from two replicates. The TGM6-D3:TβRII binding experiment was globally fit to a simple binding model from three replicates. The TGM6-D3 variant and TβRII variant binding experiments were fit to a simple binding model from 1-2 replicates per variant. Competition experiments were performed with TβRII in the syringe and the competitors in the sample cell (**Table S3**). The data were globally fit using a simple competitive binding model with one replicate per condition.

### X-ray Structure Determination

**X-** TGM6-D3 (residues 16-102 of the full-length construct) and TβRII 46-155 were mixed in a 1.1-to-1.0 ratio, with TGM6-D3 being in slight excess. The binary complex was fractionated by SEC using a HiLoad Superdex 75 26/60 column (GE Healthcare, Piscataway, NJ) in 25 mM Tris, 100 mM NaCl, 0.05% NaN_3_, pH 8.0. The fractions containing the binary complex were pooled and concentrated to 50 mg mL^-1^ for crystallization. The binary complex was crystallized in 0.1 M sodium cacodylate, 25% (w/v) PEG 4000, pH 6.5. Large star-burst-like crystal clusters with plate-like arms grew at ambient temperature in about three days.

Harvested crystals were briefly soaked in mother liquor containing 14% glycerol for cryoprotection and mounted in nylon loops with excess mother liquor wicked off. The looped crystals were then flash-cooled in liquid nitrogen prior to data collection. Data were collected at the Southeast Regional Collaborative Access Team (SER-CAT) 22-ID beamline at the Advanced Photon Source, Argonne National Laboratory and integrated and scaled using XDS [55]. The structure was determined by the molecular replacement method implemented in PHASER [56] using the 1.1 Å TβRII X-ray structure (PDB 1M9Z) [57] and the TGM1-D3 NMR structural ensemble (PDB 7SXB) [28] as search models. Coordinates were refined using REFMAC5 [58] and alternated with manual rebuilding using COOT [59]. Data collection and refinement statistics are shown in **Table S5**.

### TGF-β/TGM inhibition assays in NIH-3T3 fibroblasts

TGF-β/TGM inhibition assays utilizing NIH-3T3 cells were performed using NIH-3T3 cells stably transfected with a CAGA_12_-luciferase reporter construct as previously reported [34]. Briefly, NIH-3T3-CAGA cells were plated at a concentration of 2×10^4^ cells per well in 24 well plates containing Dulbecco’s modified Eagle’s medium (DMEM) plus 10% fetal calf serum (FCS) and allowed to attach for 18 hours. Cells were washed with PBS and incubated with DMEM plus 0.1% FCS for 6 hours. After this initial incubation, increasing concentrations of either TGM6 or TGM6-D3 were added to the wells for 30 min prior to stimulation with 0.1 ng/ml TGM1 or TGF-β (Peprotech). The cells were incubated for 15 hours and then washed with PBS and lysed using 100 μL of reporter lysis buffer (Promega). To measure luciferase activity, 30 μL Luciferase Assay Reagent (Promega) was added to 20 μL of lysate. The protein concentration of each lysate was analyzed using Bio-Rad protein assay reagent according to the manufacturer’s instructions (Biorad). Luciferase units obtained were normalized to the protein content of each well. All experiments were performed with three independent wells per condition.

### TGF-β/TGM inhibition and activation assays in MFB-F11 fibroblasts

TGF-β/TGM inhibition assays utilizing MFB-F11 cells containing a TGF-β-responsive alkaline phosphatase reporter [33] were performed as previously reported [6]. Briefly, 80-90% confluent cells were detached with trypsin, and resuspended in DMEM containing 2% FCS, 100 U mL^-1^ penicillin, 100 mg mL^-1^ streptomycin, and 2 mM L-glutamine at a concentration of 8×10^5^ cells mL^-1^. Cells were plated at 4×10^4^ (50 μL) cells per well of a 96-well flat-bottomed plate (Corning) and left to incubate at 37°C for 2 h. After this initial incubation, increasing concentrations of either full-length TGM6 or TGM6-D3 were added to the wells in a volume 25 μL. After 30 minutes, cells were stimulated with 40 or 200 pM TGF-β or TGM1 in a volume of up to 25 μL and incubated for another 24 hours at 37°C, 5% CO_2_. The final volume in each well was 100 μL. After the second incubation, 20 μL of supernatant was aspirated from each well, added to a 96-well flat-bottomed plate. Sigma FastTM p-nitrophenyl phosphate substrate was reconstituted in sterile Milli-Q water and 180 μl of it was added to each well of the 96 well plate and incubated at room temperature in the dark for up to 24h (Sigma-Aldrich). Plates were read at 405 nm on an Emax precision microplate reader (Molecular Devices). All conditions were set up in triplicate and repeated at least twice. IC_50_ values were calculated in Prism 9 (Graphpad Software, Inc.) by globally fitting the replicates of each inhibition assay to a nonlinear dose-response inhibition model.

For TGM1 activation assays, MFB-F-11 cells were seeded at 4×10^4^ (50 µL) cells per well of a 96-well flat-bottomed plate (Corning) and left to incubate at 37°C for 4 h. Increasing concentrations of full-length TGM1 orTGM1-D123 (wild type or QRRG mutant) were added to the wells to final volume of 100 µL and incubated for another twenty-four hours at 37°C, 5% CO_2_. Remainder of the assay was completed as described above.

### pSMAD stimulation and Western blotting

MFB-F11 cells were cultured in 6-well tissue culture plates (Corning) until they reached a confluency of 80-90% in complete growth medium (DMEM, 10% FBS, 1% L-glutamine, 1X penicillin-streptomycin). The growth medium was then replaced with serum-free DMEM, and the cells were incubated at 37°C with 5% CO_2_ for 4 hours. To stimulate pSMAD2, TGFβ, TGM1, or TGM1-D123 were added to the cells and incubated at 37°C for 1 hour. To inhibit pSMAD2, the cells were incubated with increasing concentrations of TGM6 for 30 minutes. Following this incubation, the cells were stimulated with either TGFβ or TGM1 for 1 hour at 37°C. The cells were washed with ice-cold PBS and lysed with RIPA buffer (0.05M Tris-HCl, pH 7.4, 0.15M NaCl, 0.25% deoxycholic acid, 1% NP-40, 1mM EDTA) containing 1X Halt protease and phosphatase inhibitors (Invitrogen). Cell lysates were cleared by centrifugation at 13000 g, 4°C for 5 minutes, and protein concentrations were estimated using the Precision Red reagent (Cytoskeleton Inc). Protein samples were prepared by mixing 1X LDS (Invitrogen), 25 mM DTT, and boiling at 100°C for 5 minutes. Equal amounts of cell lysates were analyzed on 4-12% bis-tris SDS-PAGE gels (NuPAGE, Invitrogen) and transferred onto a nitrocellulose membrane using the iBlot2 system (Invitrogen). The membranes were treated with a 5% non-fat milk blocking solution for 1 hour and incubated with the primary antibody (diluted 1:1000 in 5% BSA-containing TBST) overnight at 4°C. They were then washed three times (5 minutes each) with 1X TBST. To detect the protein bands, a fluorescently-conjugated secondary antibody (diluted 1:10000 in 5% BSA-containing TBST) was used, and the bands were visualized using the Odyssey CLx Imaging System (LI-COR Biosciences).

### Foxp3^+^ Treg induction assay

Induction of the Foxp3 transcription factor in murine splenic CD4+ T cells was conducted as previously described [60]. Briefly, a single cell suspension was prepared from the spleens of naïve Foxp3-GFP BALB/c transgenic mice [61], with 2 min incubation in red blood cell lysis buffer (Sigma), then washed and resuspended in RPMI containing HEPES (Gibco), supplemented with 2 mM L-glutamine, 100 U/ml of penicillin and 100 μg/ml of streptomycin (Gibco), 10% heat-inactivated FCS (Gibco), and 50 nM 2-mercaptoethanol (Gibco). Naive CD4+ T cells were isolated by magnetic sorting using the mouse naïve CD4+ T cell isolation kit on the AutoMACS system (Miltenyi) as per the manufacturer’s instructions. Cells were cultured at 2×10^5^ per well in flat-bottomed 96-well plates (Corning) with the addition of IL-2 (Miltenyi) at a final concentration of 400 U/ml and pre-coated with 10 µg/ml of anti-CD3 (eBioscience). Cells were cultured with TGF-β1 or TGM-1 with or without increasing concentrations of TGM6 at 37°C in 5% CO_2_ for 72 h before being removed for flow cytometric analysis. Briefly, cells were washed with PBS and stained with eBioscience Fixable Viability Dye eFluor 506 (Invitrogen) and TruStain FcX PLUS (anti-mouse CD16/32) Antibody (Biolegend) to identify live cells and prevent unspecific binding, respectively. Following this, cells were incubated with fluorochrome-conjugated anti-CD4 (BV650, Biolegend). Cells were analyzed using a BD FACSCelesta Cell Analyzer (BD) and data analyzed by FlowJo software (BD). Analysis of stained cells was performed on single live cells.

### Transcriptional Florescent Protein–Based Reporter Assays

The CAGA-MLP-dynGFP lentiviral vector was used as previously described [62]. The BRE-MLP-mCHERRY-d2 lentiviral reporter was made using the pGL3-MLP-BRE-Luc plasmid (cloning information available on request) [63]. Lentiviruses were generated by transfection HEK293T cells with packaging constructs and the lentiviral constructs using standard protocols. Cells were exposed to lentivirus containing pLV-CAGA-MLP-dynGFP or pLV-BRE-MLP-mCHERRYd2 for 48 h, after which the cells were selected using puromycin (CAGA-MLP-dynGFP) or blasticidin (BRE-MLP-mCHERRYd2). Murine NIH-3T3 fibroblasts containing either the CAGA-MLP-dynGFP or the BRE-MLP-mCHERRYd2 reporter, or murine NM-18 epithelial cells containing the CAGA-MLP-dynGFP reporter, were seeded in 96-well plates. For the CAGA-MLP-dynGFP reporter, the cells were stimulated the next day with either TGF-β3 or activin A in 10% serum. For the BRE-MLP-mCHERRYd2 the cells were put on media without serum overnight and were subsequently stimulated with BMPs. Directly after ligand stimulation the cells were placed in the IncuCyte S3 live-cell imaging analysis system (Sartorius). The cells were imaged every 3 h for a period of 48 h. Fluorescence intensity was analyzed using the IncuCyte software.

### Statistical analyses

Statistical analyses were performed using Prism 9 (Graphpad Software, Inc.) or Sedphat [53, 54], as appropriate. For comparisons of two groups, a Student’s unpaired two-tailed t-test was used, assuming unequal variance. *P* values of < 0.05 were considered statistically significant. Sample sizes were chosen empirically based on the laboratory’s previous experience in the calculation of experimental variability; sample sizes for each experiment were not pre-determined by individual power calculations.

## ACKNOWLEDGEMENTS

We would like to thank Chang Byeon for his assistance in producing and purifying proteins used in the binding studies and Matthew Whitley for his assistance with analysis of the diffraction data. This research was supported by the NIH through an RO1 (GM58670) and RO3 (AI53915) awarded to A.H and an F30 (AI157069) awarded to A.M. and the Wellcome Trust through an Investigator Award (Ref 219530) awarded to R.M.M. and the Wellcome Trust core-funded Wellcome Centre for Investigative Parasitology (Ref 104111). X-ray data were collected at Southeast Regional Collaborative Access Team (SER-CAT) 22-ID beamline at the Advanced Photon Source, Argonne National Laboratory. SER-CAT is supported by its member institutions, and equipment grants (RR25528, RR028976 and OD027000) from the NIH. Molecular graphics and analyses performed with UCSF ChimeraX, developed by the Resource for Biocomputing, Visualization, and Informatics at the University of California, San Francisco, with support from NIH GM129325 and the Office of Cyber Infrastructure and Computational Biology, National Institute of Allergy and Infectious Diseases.

## ABBREVIATIONS

ActRII: Activin receptor type II
ActRIIb: Activin receptor type IIb
ALK1: Activin receptor-like kinase 1
ALK2: Activin receptor-like kinase 2
ALK3: Activin receptor-like kinase 3, Bone morphogenic protein receptor, type 1A
ALK4: Activin receptor-like kinase 4, Activin receptor type-1B
ALK5: Activin receptor-like kinase 5, TGF-βtype I receptor
BMP: Bone morphogenic protein
BMPRII: BMP type II receptor
BSA: Bovine serum albumin
CCP: complement control protein
DMEM: Dulbecco’s modified Eagle’s medium
FBS: Fetal bovine serum
TGM6: full-length TGM6
HSQC: heteronuclear single quantum correlation
ITC: isothermal titration calorimetry
NMR: nuclear magnetic resonance
TGF-β: transforming growth factor beta
TGM: TGF-β mimic
TGM1: TGF-β mimic 1
TGM1-D123: TGF-β mimic 1 domains 1, 2, and 3
TGM1-D3: TGF-β mimic 1 domain 3
TGM6: TGF-β mimic 6
TGM6-D3: TGM6 domain 3
TGM6-D45: TGM6 domains 4 and 5
Treg: regulatory T-cell
TβRII: TGF-β type II receptor

## Notes

### Competing Interest Statement

The authors have declared no competing interest.

