## Supplementary Information for "TGM6, a helminth secretory product, mimics TGF-β binding to TβRII to antagonize TGF-β signaling in fibroblasts"

**Supporting Information**

### TGM6, a secretory product of the helminth *H. polygyrus*, mimics TGF-β binding to TβRII to block signaling in fibroblasts

Stephen E. White, Tristin A. Schwartze, Ananya Mukundan, Christina Schoenherr, Shashi P. Singh, Maarten van Dinther , Kyle T. Cunningham, Madeleine P. J. White, Tiffany Campion, John Pritchard, Cynthia S. Hinck, Peter ten Dijke, Gareth Inman, Rick M. Maizels, and Andrew P. Hinck

10 Tables

12 Figures

**Table S1. TGM6:TβRII binding as assessed by ITC**

| | | |
|---|---|---|
| Cell | 32.3 μM TGM6 | 14.7 μM TGM6-D3 |
| Syringe | 339.7 μM TβRII | 200.0 μM TβRII |
| Cell concentration (μM) | 32.3 | 14.7 |
| Temperature (°C) | 25 | 35 |
| N (sites) | 1.16 | 1.01 |
| $K_D$ (nM) [a] | 222 ± 97.1 [b,c] | 437 ± 81 [b,d] |
| ΔH (kcal $mol^{-1}$) [a] | -8.918 ± 0.231 [b] | -21.268 ± 0.651 [b] |
| ΔG (kcal $mol^{-1}$) [a] | -9.066 | -8.967 |
| -TΔS (kcal $mol^{-1}$) [a] | 0.148 | 12.301 |

[a] Number of sites set to “1” for analysis.
[b] Uncertainty reported as ± 1$\sigma$.
[c] Global fit of two replicates.
[d] Global fit of three replicates.

**Table S2. Type II receptor binding to TGM6-D3 as assessed by ITC**

| | | | | | | |
|---|---|---|---|---|---|---|
| Cell | TGM6-D3 | Buffer | TGM6-D3 | Buffer | TGM6-D3 | Buffer |
| Syringe | ActRII | ActRII | ActRIIb | ActRIIb | BMPRII | BMPRII |
| Cell concentration (μM) | 7.35 | 0.00 | 10.0 | 0.00 | 20.0 | 0.00 |
| Syringe concentration (μM) | 60.5 | 60.5 | 100 | 100 | 215 | 215 |
| Temperature (°C) | 35 | 35 | 25 | 25 | 35 | 35 |
| N (sites) | ND[a] | ND[a] | ND[a] | ND[a] | ND[a] | ND[a] |
| $K_D$ (nM) | ND[a] | ND[a] | ND[a] | ND[a] | ND[a] | ND[a] |
| ΔH (kcal $mol^{-1}$) | ND[a] | ND[a] | ND[a] | ND[a] | ND[a] | ND[a] |
| ΔG (kcal $mol^{-1}$) | ND[a] | ND[a] | ND[a] | ND[a] | ND[a] | ND[a] |
| -TΔS (kcal $mol^{-1}$) | ND[a] | ND[a] | ND[a] | ND[a] | ND[a] | ND[a] |

[a] Not determined due to weak or no signal

**Table S3. TβRII competition binding as assessed by ITC**

| | | |
|---|---|---|
| Cell | 15 μM TGF-β2-7M2R | 15 μM TGM6-D3 |
| Syringe | 105 μM TβRII | 150 μM TβRII |
| Competitor [a] | 0.0 or 6.0 μM TGM6-D3 | 0.0 or 6.0 μM TGM1-D3 |
| Temperature (°C) | 35 | 25 |
| $K_D$ (nM) | 19.34 (7.81, 39.57)[b] | 188.6 (108.8, 308.0)[c] |
| ΔH (kcal mol$^{-1}$) | -14.240 (-14.8531, -13.6366)[b] | -10.382 (-10.9570, -9.8751)[c] |
| ΔG (kcal mol$^{-1}$) | -10.876[d] | -9.174[e] |
| -TΔS (kcal mol$^{-1}$) | 3.364[d] | 1.208[e] |

[a] Competitor was added to the sample cell.
[b] $K_D$ and ΔH correspond to the parameters, derived from the global fit, for TβRII:mmTGF-β2-7M2R binding in the absence of competitor; uncertainty is reported as the limits of the ±1$\sigma$ confidence interval.
[c] $K_D$ and ΔH correspond to the parameters, derived from the global fit, for TGM6-D3:TβRII binding in the absence of competitor; uncertainty is reported as the limits of the ±1$\sigma$ confidence interval.
[d] ΔG and -TΔS correspond to those for TβRII:mmTGF-β2-7M binding in the absence of competitor calculated from ΔG = ΔH – TΔS and globally fitted values for $K_D$ and ΔH.
[e] ΔG and -TΔS correspond to those for TβRII:TGM6-D3 binding in the absence of competitor calculated from ΔG = ΔH – TΔS and globally fitted values for $K_D$ and ΔH.

**Table S4. TGM1-D45:mCD44 and TGM6-D45:mCD44 binding as assessed by ITC**

| Syringe | Cell | Temp (°C) | N (sites) | $K_D$ (nM) | ΔH (kcal mol$^{-1}$) | ΔG (kcal mol$^{-1}$) | -TΔS (kcal mol$^{-1}$) |
|---|---|---|---|---|---|---|---|
| 90 μM mCD44 | 8 μM TGM6-D45 | 35 | ND | ND | ND | ND | ND |
| 100 μM mCD44 | 15 μM TGM1-D45 | 35 | 0.64[a] | 56 (28, 106)[b] | -19.5 (-20.9, -18.5)[b] | -10.2 | 9.3 |

[a] Number of sites determined by incompetent fraction value on Sedphat; set to '1' for thermodynamic analysis.
[b] Uncertainty reported as 68.3% confidence interval.

**Table S5. Crystallographic data, phasing, and refinement of the TGM6-D3:TβRII complex**

| | |
|---|---|
| **Data Collection** | |
| X-ray Source | APS BEAMLINE 22-ID |
| Wavelength | 1.00 Å |
| Detector | DECTRIS EIGER X 16M |
| **Data Reduction** | |
| Space Group | $P2_12_12$ |
| a,b,c (Å) | 56.04, 128.39, 29.65 |
| α,β,γ (°) | 90, 90, 90 |
| Completeness Overall (%) | 98.5 (85.1) |
| Resolution (Å) | 42.80-1.40 (7.67-1.40) |
| $R_{meas}$ | 0.079 (1.428) |
| $R_{pim}$ | 0.022 (0.506) |
| <I/σ(I)> | 17.5 (1.6) |
| Redundancy or Multiplicity | 12.4 (7.5) |
| $CC_{1/2}$ | 0.999 (0.700) |
| CC* | 1.000 (0.907) |
| Matthews Coefficient | 2.34 |
| No. Observations | 528172 |
| No. Reflections | 42468 |
| **Refinement** | |
| 1:1 complexes in the asymmetric unit | 1 |
| Resolution (Å) | 42.253 - 1.401 |
| $R_{work}$ / $R_{free}$ | 0.2180 / 0.2308 |
| Residues | 105, 80 |
| Atoms / Non-Hydrogens | 3129 / 1669 |
| Protein Overall / Heavy / Backbone | 2931 / 1505 / 745 |
| Ion and Ligand Overall / Heavy | 63 / 29 |
| Water(s) | 135 |
| B-factors | 28.7 |
| Protein Overall / Heavy / Backbone | 28.38 / 27.7 / 24.8 |
| Ion and Ligand Overall / Heavy | 25.58 / 27.28 |
| Water(s) | 37.13 |
| R.M.S. Deviations | |
| Bond-Lengths | 0.0127 |
| Bond-Angles | 1.6576 |
| MolProbity | |
| Clash Score | 0 |
| MolProbity Score | 0.65 |
| Ramachandran | |
| Allowed | 100% |
| Favored | 97.18% |
| Outliers | 0% |
| Rotamers | |
| Allowed | 100% |
| Favored | 94.89 % |
| Outliers | 0% |
| Clashes | 0 |
| **Deposition** | |
| RCSB PDB | 8GDT |

[a] The numbers in the Data Reduction section correspond to the value for the overall and in parentheses the high resolution shell.

#### Table S6. TGM6-D3 WT:TβRII variant and TβRII WT:TGM6-D3 variant binding as assessed by ITC at 25 °C

| Syringe | Cell | Temp (°C) | N (sites) | $K_D$ (μM) | $\Delta H$ (kcal $mol^{-1}$) | $\Delta G$ (kcal $mol^{-1}$) | $-T\Delta S$ (kcal $mol^{-1}$) |
|---|---|---|---|---|---|---|---|
| 426 μM TGM6-D3 | 25 μM TβRII WT | 25 | 0.68[a] | 0.35 (0.28, 0.43)[b] | -16.3 (-16.8, -15.9)[b] | -8.8 | 7.5 |
| 590 μM TGM6-D3 | 30 μM TβRII D55A | 35 | 0.52 | 2.25 (2.05, 2.45)[b] | -22.1 (-22.6, -21.5)[b] | -8.0 | 14.1 |
| 290 μM[a] TGM6-D3 | 18.4 μM TβRII I76A | 25 | 0.69[a] | 7.35 (6.40, 8.51)[b] | -10.6 (-11.4, -10.0)[b] | -7.0 | 3.6 |
| 290 μM[a] TGM6-D3 | 18.9 μM TβRII D141A | 25 | 0.75[a] | 0.94 (0.87 1.00)[b] | -11.5 (-11.7, -11.4)[b] | -8.2 | 3.3 |
| 590 μM[a] TGM6-D3 | 30 μM TβRII E142A | 35 | 0.56 | 3.17 (2.75, 3.67)[b] | -21.8 (-23.1, -20.8)[b] | -7.8 | 14.1 |
| 300 μM TβRII | 10 μM TGM6-D3 WT | 25 | 0.87[a] | 0.36 (0.31, 0.41)[b] | -11.3 (-11.6, -11.1)[b] | -8.8 | 2.5 |
| 667 μM TβRII | 25 μM TGM6-D3 R38A | 25 | 0.85[a] | 8.03 (7.61, 8.48)[b] | -17.0 (-17.4, -16.6)[b] | -7.0 | 10.0 |
| 667 μM TβRII | 25 μM TGM6-D3 I78A | 25 | 0.71[a] | 5.56 (5.24, 5.71)[b] | -17.5 (-17.6, -17.3)[b] | -7.2 | 10.3 |
| 667 μM TβRII | 25 μM TGM6-D3 Y80A | 25 | 0.33[a] | 48.4 (38.1, 61.8)[b] | -21.5 (-26.6, -17.7)[b] | -5.9 | 15.6 |
| 150 μM TβRII | 10 μM TGM6-D3 Y80F | 25 | 0.67[a] | 1.25 (1.04, 1.49)[b] | -9.6 (-10.2, -9.1)[b] | -8.1 | 1.6 |
| 200 μM TβRII | 10 μM TGM6-D3 R82A | 35 | 0.93 | 4.23 (3.45, 5.27)[b] | -16.6 (-18.9, -15.0)[b] | -7.6 | 9.1 |
| 200 μM TβRII | 10 μM TGM6-D3 R82S | 35 | 0.95 | 2.65 (2.37, 4.11)[b] | -19.4 (-18.8, -15.1)[b] | -7.9 | 11.6 |
| 667 μM TβRII | 25 μM TGM6-D3 Y93A | 25 | 0.17[a] | 39.0 (30.4, 50.8)[b] | -19.2 (-23.2, -16.3)[b] | -6.0 | 13.2 |
| 150 μM TβRII | 10 μM TGM6-D3 R95A | 25 | 1.00[a] | 3.82 (1.50, 15.18)[b] | -13.3 (-30.3, -9.4)[b] | -7.4 | 5.9 |
| 150 μM TβRII | 10 μM TGM6-D3 P94K R95N (KN) | 25 | 0.90[a] | 0.27 (0.23, 0.30)[b] | -13.4 (-13.6, -13.1)[b] | -9.0 | 4.4 |
| 120 μM TβRII | 10 μM TGM6-D3 Q81K R82S R83G G84T (KSGT) | 35 | 0.82 | 4.49 (1.74, 9.76)[c] | -21.6 (-39.8, -14.2)[c] | -7.5 | 14.0 |
| 100 μM TβRII | 10 μM TGM1-D3 | 35 | 1.07 | 1.53 (0.68, 4.48) | -9.3 (-15.2, -7.2) | -8.2 | 1.1 |
| 360 μM TβRII | 30 μM TGM1-D3 S242R | 35 | 0.70 | 0.38 (0.24, 0.56) | -11.6 (-12.4, -10.9) | -9.1 | 2.5 |
| 120 μM TβRII | 10 μM TGM1-D3 K254P N255R (PR) | 35 | 0.91 | 2.62 (0.39, 28.20)[d] | -5.2 (-9.3, -3.4)[d] | -7.9 | -2.6 |
| 120 μM TβRII | 10 μM TGM1-D3 K241Q S242R G243R T244G (QRRG) | 35 | 1.10 | 0.18 (0.07, 0.40) | -8.4 (-9.5, -7.4) | -9.5 | -1.13 |

[a] Number of sites determined by incompetent fraction value on Sedphat; set to '1' for thermodynamic analysis.
[b] Uncertainty reported as 68.3% confidence interval.
[c] Fit by constraining either the $\Delta H$ to -31.5, -11.5 or the $K_D^{-1}$ to (5.05, 5.65) x $10^5$
[d] Fit by constraining either the $\Delta H$ to -15.2, -0.2 or the $K_D^{-1}$ to (5.28, 5.88) x $10^5$

**Table S7. *H. polygyrus* TGM constructs used in this study**

<table>
<tr><th>Construct</th><th>Residue range and features*</th><th>Sequence</th></tr>
<tr><td>TGM6</td><td>Residues 17-254 of H. polygyrus TGF-β Mimic 6 (NCBI MG429741)<br><br>Expressed as Igk Signal Peptide-TGM6-Linker-Myc Tag-Linker-His6 fusion</td><td>METDTLLLWV LLLWVPGSTG DAAQPARRAS CPPLPDDETV<br>WYEYYGYVDG RHTVGDAAIK DSLENYPPNT HARRHCKALS<br>KKADPGEFVA ICYQRRGTSE SQWQYYPRIA SCPDPRCKPL<br>EKNDSVSYEY FTKPTKGLKM GSITKPDKSG KYPEETFVRR<br>YCNDLPRNSL AQGKTYAECL DSEWKLKNLP DCRFAAGCDE<br>EYLLEKLMFV DISYWGKDAA KFSDDKTYRY YRPGSKVTAK<br>CKGKSVKLTC VDGGYWVTVD GRKALCTAAA RGGPEQKLIS<br>EEDLNSAVDH HHHHH</td></tr>
<tr><td>TGM6-D3</td><td>Residues 15-102 of H. polygyrus TGF-β Mimic 6 (NCBI MG429741)<br><br>Expressed as a Thioredoxin- His10-Linker-Thrombin-Linker-TGM6-D3 fusion</td><td>MSDKIIHLTD DSFDTDVLKA DGAILVDFWA EWCGPCKMIA<br>PILDEIADEY QGKLTVAKLN IDQNPGTAPK YGIRGIPTLL<br>LFKNGEVAAT KVGALSKGQL KEFLDANLAG SGSGHMSSGH<br>HHHHHHHHHS SGGSGLVPR|G SGTGSSCPPL PDDETVWYEY<br>YGYVDGRHTV GDAAIKDSLE NYPPNTHARR HCKALSKKAD<br>PGEFVAICYQ RRGTSESQWQ YYPRIASCPD P</td></tr>
<tr><td>TGM6-D45</td><td>Residues 103-254 of H. polygyrus TGF-β Mimic 6 (NCBI MG429741)<br><br>Expressed as a Thioredoxin- His10-Linker-Thrombin-Linker-TGM-D45 fusion</td><td>MSDKIIHLTD DSFDTDVLKA DGAILVDFWA EWCGPCKMIA<br>PILDEIADEY QGKLTVAKLN IDQNPGTAPK YGIRGIPTLL<br>LFKNGEVAAT KVGALSKGQL KEFLDANLAG SGSGHMSSGH<br>HHHHHHHHHS SGGSGLVPR|G SGTRCKPLEK NDSVSYEYFT<br>KPTKGLKMGS ITKPDKSGKY PEETFVRRYC NDLPRNSLAQ<br>GKTYAECLDS EWKLKNLPDC RFAAGCDEEY LLEKLMFVDI<br>SYWGKDAAKF SDDKTYRYYR PGSKVTAKCK GKSVKLTCVD<br>GGYWVTVDGR KALCT</td></tr>
<tr><td>TGM1</td><td>Residues 16-422 of H. Polygyrus TGF-β Mimic 1 (NCBI ATO59092.1)<br><br>Expressed as Igk Signal Peptide-Linker-TGM1-Linker-Myc Tag-Linker-His6 fusion</td><td>METDTLLLWVLLLWVPGSTGDAAQPARRADDSGCMPFSDEAAT<br>YKYVAKGPKNIEIPAQIDNSGMYPDYTHVKRFCKGLHGEDTTG<br>WFVGICLASQWYYYEGVQECDDRRCSPLPTNDTVSFEYLKATV<br>NPGIIFNITVHPDASGKYPELTYIKRICKNFPTDSNVQGHIIG<br>MCYNAEWQFSSTPTCPASGCPPLPDDGIVFYEYYGYAGDRHTV<br>GPVVTKDSSGNYPSPTHARRRCRALSQEADPGEFVAICYKSGT<br>TGESHWEYYKNIGKCPDPRCKPLEANESVHYEYFTMTNETDKK<br>KGPPAKVGKSGKYPEHTCVKKVCSKWPYTCSTGGPIFGECIGA<br>TWNFTALMECINARGCSSDDLFDKLGFEKVIVRKGEGSDSYKD<br>DFARFYATGSKVIAECGGKTVRLECSNGEWHEPGTKTVHRCTK<br>DGIRTLGPEQKLISEEDLNSAVDHHHHHH-</td></tr>
<tr><td>TGM1-D123</td><td>Residues 16-262 of H. Polygyrus TGF-β Mimic 1 (NCBI ATO59092.1)<br>Expressed as Igk Signal Peptide-Linker-TGM1-D123-Linker-Myc Tag-Linker-His6 fusion</td><td>METDTLLLWVLLLWVPGSTGDAAQPARRADDSGCMPFSDEAAT<br>YKYVAKGPKNIEIPAQIDNSGMYPDYTHVKRFCKGLHGEDTTG<br>WFVGICLASQWYYYEGVQECDDRRCSPLPTNDTVSFEYLKATV<br>NPGIIFNITVHPDASGKYPELTYIKRICKNFPTDSNVQGHIIG<br>MCYNAEWQFSSTPTCPASGCPPLPDDGIVFYEYYGYAGDRHTV<br>GPVVTKDSSGNYPSPTHARRRCRALSQEADPGEFVAICYKSGT<br>TGESHWEYYKNIGKCPDPGPEQKLISEEDLNSAVDHHHHHH-</td></tr>
</table>

| | | |
|---|---|---|
| TGM1-D3 | Residues 177-262 of *H. Polygyrus* TGF-β Mimic 1 (NCBI ATO59092.1)<br><br>Expressed as a Thioredoxin- His6-Linker-Thrombin-Linker-TGM-D3 fusion | MSDKIIHLTD DSFDTDVLKA DGAILVDFWA EWCGPCKMIA PILDEIADEY QGKLTVAKLN IDQNPGTAPK YGIRGIPTLL LFKNGEVAAT KVGALSKGQL KEFLDANLAG SGSGHMHHHH HHSSGLVPR\|G SGTGCPPLPD DGIVFYEYYG YAGDRHTVGP VVTKDSSGNY PSPTHARRRC RALSQEADPG EFVAICYKSG TTGESHWEYY KNIGKCPDP |

*All residue numbering begins with the N-terminal methionine of the naturally occurring signal peptide

**Table S8. Type I receptor constructs used in this study**

| Construct | Residue range and features* | Sequence |
|---|---|---|
| ALK1 (TSRI) | Residues 22-118 of human ALK1 (TSR1) (NCBI NP_000011)<br><br>Expressed as a Linker-His6-Linker-Thrombin-Linker-Alk1 fusion | MGSSHHHHHH SSGLVPR\|GSH MDPVKPSRGP LVTCTCESPH CKGPTCRGAW CTVVLVREEG RHPQEHRGCG NLHRELCRGR PTEFVNHYCC DSHLCNHNVS LVLEATQPPS EQPGTDGQ |
| ALK2 (ActRIA) | Residues 21-120 of human ALK2 (ActRIA) (NCBI NP_001096)<br><br>Expressed as a Thioredoxin-His6-Linker-Thrombin-Linker-Alk2 fusion | MSDKIIHLTD DSFDTDVLKA DGAILVDFWA EWCGPCKMIA PILDEIADEY QGKLTVAKLN IDQNPGTAPK YGIRGIPTLL LFKNGEVAAT KVGALSKGQL KEFLDANLAG SGSGHMHHHH HHSSGLVPR\|G SGTMEDEKPK VNPKLYMCVC EGLSCGNEDH CEGQQCFSSL SINDGFHVYQ KGCFQVYEQG KMTCKTPPSP GQAVECCQGD WCNRNITAQL PTKGKSFPGT QNF |
| ALK3 (BMPRIA) | Residues 24-152 of human ALK3 (BMPRIA) (NCBI NP_001393488)<br><br>Expressed as a Thioredoxin-His6-Linker-Thrombin-Linker-Alk3 fusion | MSDKIIHLTD DSFDTDVLKA DGAILVDFWA EWCGPCKMIA PILDEIADEY QGKLTVAKLN IDQNPGTAPK YGIRGIPTLL LFKNGEVAAT KVGALSKGQL KEFLDANLAG SGSGHMHHHH HHSSGLVPR\|G SGTQNLDSML HGTGMKSDSD QKKSENGVTL APEDTLPFLK CYCSGHCPDD AINNTCITNG HCFAIIEEDD QGETTLASGC MKYEGSDFQC KDSPKAQLRR TIECCRTNLC NQYLQPTLPP VVIGPFFDGS IR |
| ALK4 (ActRIB) | Residues 29-107 of human ALK4 (ActRIB) (NCBI NP_001399711)<br><br>Expressed as a Linker-His6-Linker-Thrombin-Linker-Alk4 fusion | MGSSHHHHHH SSGLVPR\|GSH MVQALLCACT SCLQANYTCE TDGACMVSIF NLDGMEHHVR TCIPKVELVP AGKPFYCLSS EDLRNTHCCY TDYCNRIDLR |
| ALK5 (TβRI) | Residues 25-125 of human ALK5 (TβRI) (NCBI NP_004603.1)<br><br>Expressed as Linker-His6-Linker-Thrombin-Linker-TbRI fusion | MGSSHHHHHH SSGLVPR\|GSH MAALLPGATA LQCFCHLCTK DNFTCVTDGL CFVSVTETTD KVIHNSSCIA EIDLIPRDRP FVCAPSSKTG SVTTTYCCNQ DHCNKIELPT TVKSSPGLGP VE |

*All residue numbering begins with the N-terminal methionine of the naturally occurring signal peptide

**Table S9. Type II receptor constructs used in this study**

| Construct | Residue range and features* | Sequence |
|---|---|---|
| ActRII | Residues 20-121 of the human Activin type II receptor (NCBI NP_001265508)<br><br>Expressed as Linker-His6-Linker-Thrombin-Linker-ActRII fusion | MGSSHHHHHH SSGLVPR\|GSH MAILGRSETQ ECLFFNANWE KDRTNQTGVE PCYGDKDKRR HCFATWKNIS GSIEIVKQGC WLDDINCYDR TDCVEKKDSP EVYFCCCEGN MCNEKFSYFP EME |
| ActRIIb | Residues 25-117 of the human Activin type IIb receptor (NCBI NP_001097)<br><br>Expressed as Linker-His6-Linker-Thrombin-Artifact-Linker-ActRIIb fusion | MGSSHHHHHH SSGLVPR\|GSH MLEDPVPETR ECIYYNANWE LERTNQSGLE RCEGEQDKRL HCYASWRNSS GTIELVKKGC WLDDFNCYDR QECVATEENP QVYFCCCEGN FCNERFTHLP |
| BMPRII | Residues 29-133 of the human BMP type II receptor (NCBI NP_001195)<br><br>Expressed as Signal-Linker-His6-Linker-Thrombin-Linker-BMPRII fusion | MKWVTFLLLL FISGSAFSAA AGSSHHHHHH SSGLVPR\|GSH MNQERLCAFK DPYQQDLGIG ESRISHENGT ILCSKGSTCY GLWEKSKGDI NLVKQGCWSH IGDPQECHYE ECVVTTTPPS IQNGTYRFCC CSTDLCNVNF TENFPP |
| TβRII | Residues 38-153 of the human TGF-β type II receptor (NCBI NP_003233)<br><br>Expressed as TβRII alone, with no tags or otherwise | MVTDNNGAVK FPQLCKFCDV RFSTCDQKSC MSNCSITSIC EKPQEVCVAV WRKNENITLE TVCHDPKLPY HDFILEDAAS PKCIMKEKKK PGETFFMCSC SSDECNDNII FSEEY |

*All residue numbering begins with the N-terminal methionine of the naturally occurring signal peptide

**Table S10. Growth factor constructs used in this study**

| Construct | Residue range and features* | Sequence |
|---|---|---|
| mmTGF-β2-7M2R | Residues 303-352 and 377-414 of mouse TGF-β2 (NCBI NP_0033393) connected by an engineered loop<br><br>C379R substitution renders the protein monomeric; K327R, R328K, V381R, L391V, I394V, K396R, T397K, and I400V substitutions enable high affinity TβRII binding and high solubility | ALDAAYCFRN VQDNCCLRPL YIDFRKDLGW KWIHEPKGYN ANFCAGACPY RASKSPRCRS QDLEPLTIVY YVGRKPKVEQ LSNMIVKSCK CS |
| mCD44 | Residues 23 -174 of mouse CD44 (NCBI XP_006498709)<br><br>Expressed as Signal-His6-Linker-Thrombin-mCD44 | MKWVTFLLLL FISGSAFSGS HHHHHHGSLV PRGSHQQIDL NVTCRYAGVF HVEKNGRYSI SRTEAADLCQ AFNSTLPTMD QMKLALSKGF ETCRYGFIEG NVVIPRIHPN AICAANHTGV YILVTSNTSH YDTYCFNASA PPEEDCTSVT DLPNSFDGPV TITIVNRDGT RYSKKGEYRT HQEDID |

*All residue numbering begins with the N-terminal methionine of the naturally occurring signal peptide

**Figure S1.** TGM6-D3 and -D45 are expressed as natively-folded proteins. $^{1}H$-$^{15}N$ HSQC spectra of (A) TGM6-D3 and (B) TGM6-D45 show well-dispersed peaks outside of the random coil region (7.8 – 8.6 ppm $^{1}H$).

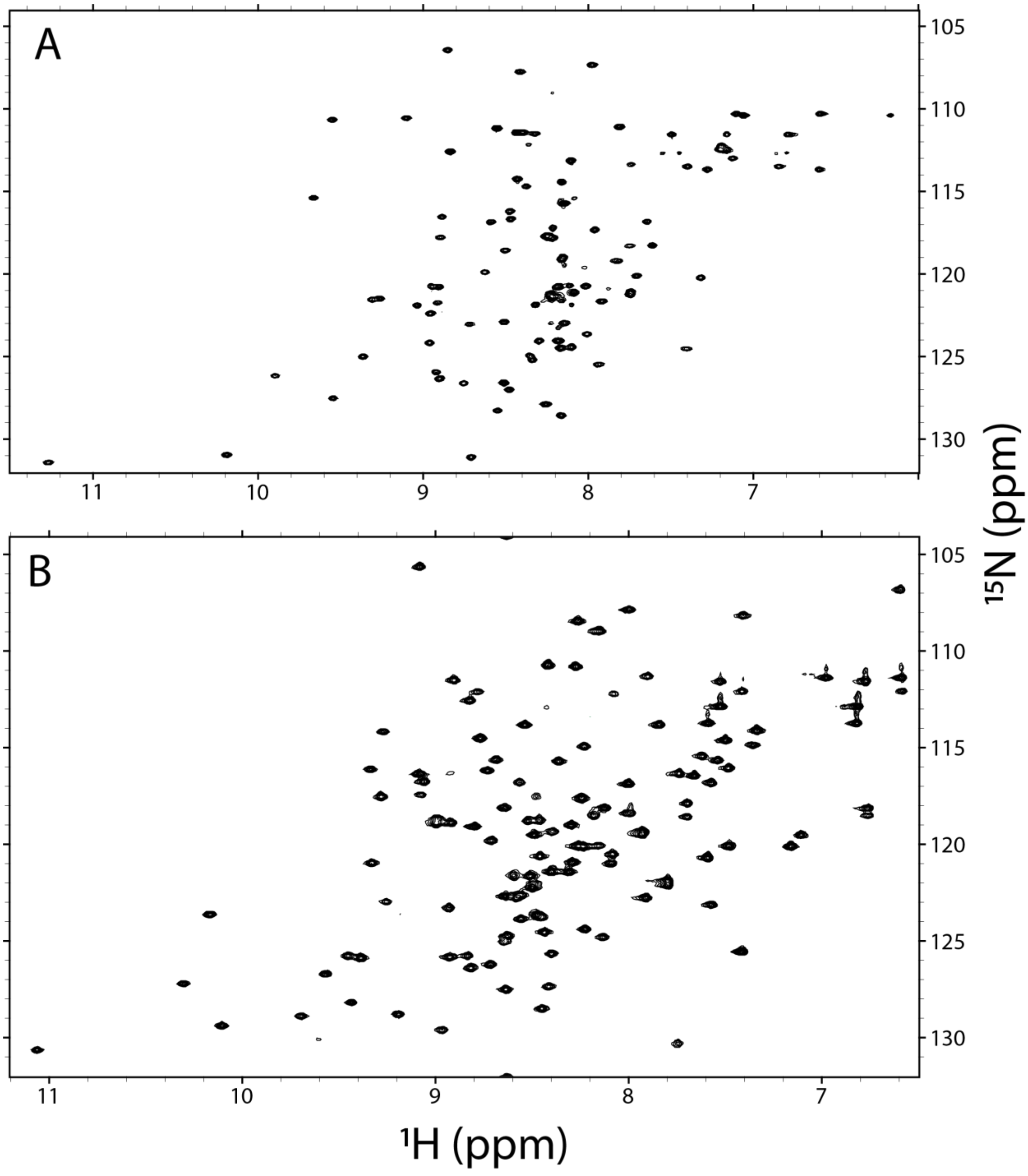

**Figure S2. TGM6-D3 does not bind ActRII, ActRIIb, or BMPRII.** Thermograms obtained upon the injection of ActRII, ActRIIb, or BMPRII into TGM6-D3 or Buffer. Panels A, C, and E correspond to the injection of ActRII, ActRIIb, and BMPRII into TGM6-D3, respectively; panels B, D, and F correspond to the injection of ActRII, ActRIIb, and BMPRII into buffer, respectively.

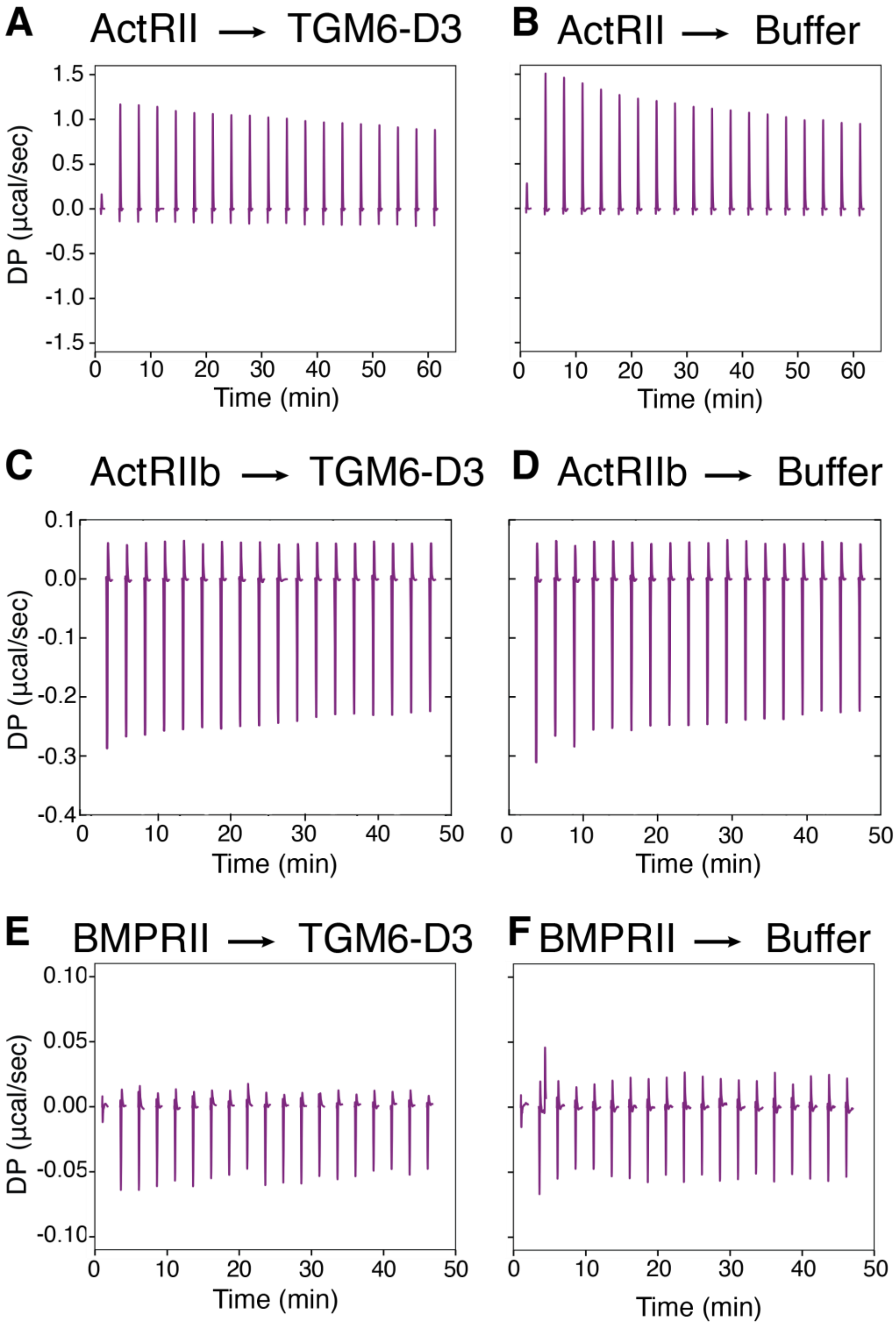

**Figure S3. TGM6-D45 does not bind to any type I receptors.** $^{1}H$-$^{15}N$ HSQC spectra of $^{15}N$-labeled type I receptors bound to an excess of unlabeled TGM6-D45 (red) overlaid onto the spectra of the type I receptors alone (blue). The receptors tested were: (A) $^{15}N$-ALK1; (B) $^{15}N$-ALK2; (C) $^{15}N$-ALK3; and (D) $^{15}N$-ALK4.

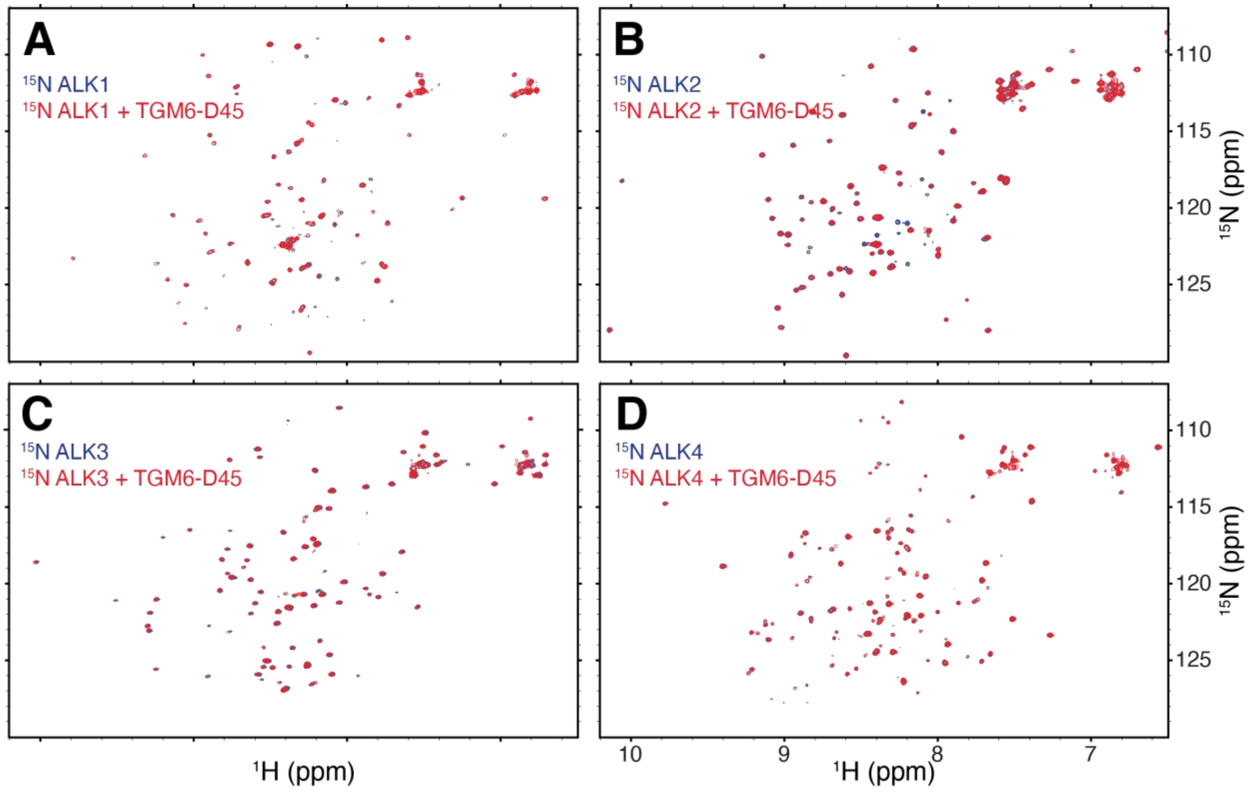

**Figure S4. The TGM6:TβRII binary complex does not bind to any type I receptors.** $^{1}H$-$^{15}N$ HSQC spectra of $^{15}N$-labeled type I receptors as bound to an excess of unlabeled TGM6:TβRII binary complex (red) overlaid onto the spectra of the type I receptors alone (blue). The receptors tested were: (A) $^{15}N$-ALK1; (B) $^{15}N$-ALK2; (C) $^{15}N$-ALK3; and (D) $^{15}N$-ALK4.

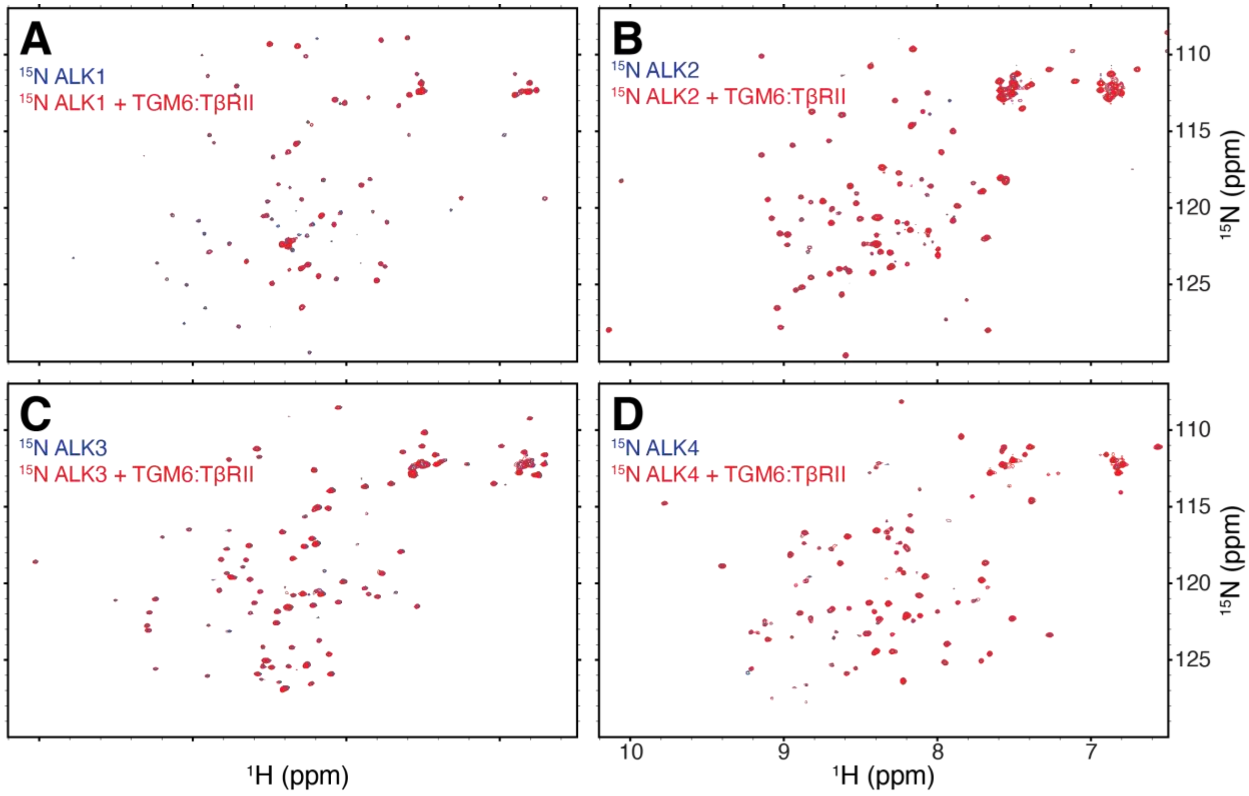

**Figure S5.** 1D $^{1}H$ NMR spectra (**A, C, E**) and mass spectra (**B, D, F**) confirming the identity and native folding of (**A, B**) ActRII, (**C, D**) ActRIIb, and (**E, F**) BMPRII used in the ITC experiments.

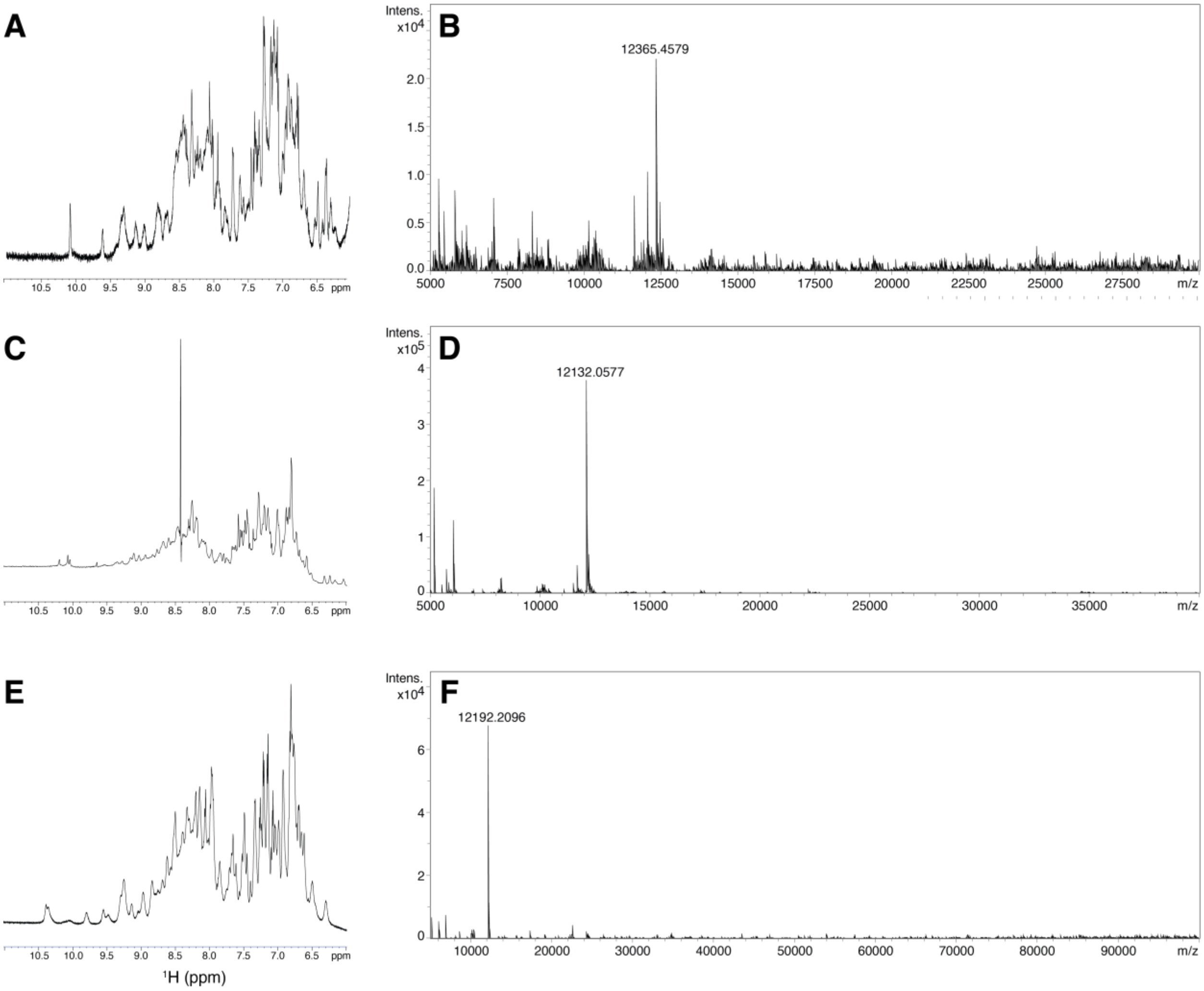

**Figure S6. TGM6 does not stimulate BMP activity nor does it inhibit BMP or activin signaling activity. A.** Stimulation of BMP mCherry reporter in NIH3T3 cells by BMP2, BMP6, and BMP7, and TGM6 at the concentrations shown in either the absence or presence of 3.6 nM TGM-6. **B.** Stimulation of TGF-β/Activin GFP reporter in NIH3T3 cells by activinA or GDF8? at the concentrations shown in either the absence or presence of 3.6 nM TGM-6.

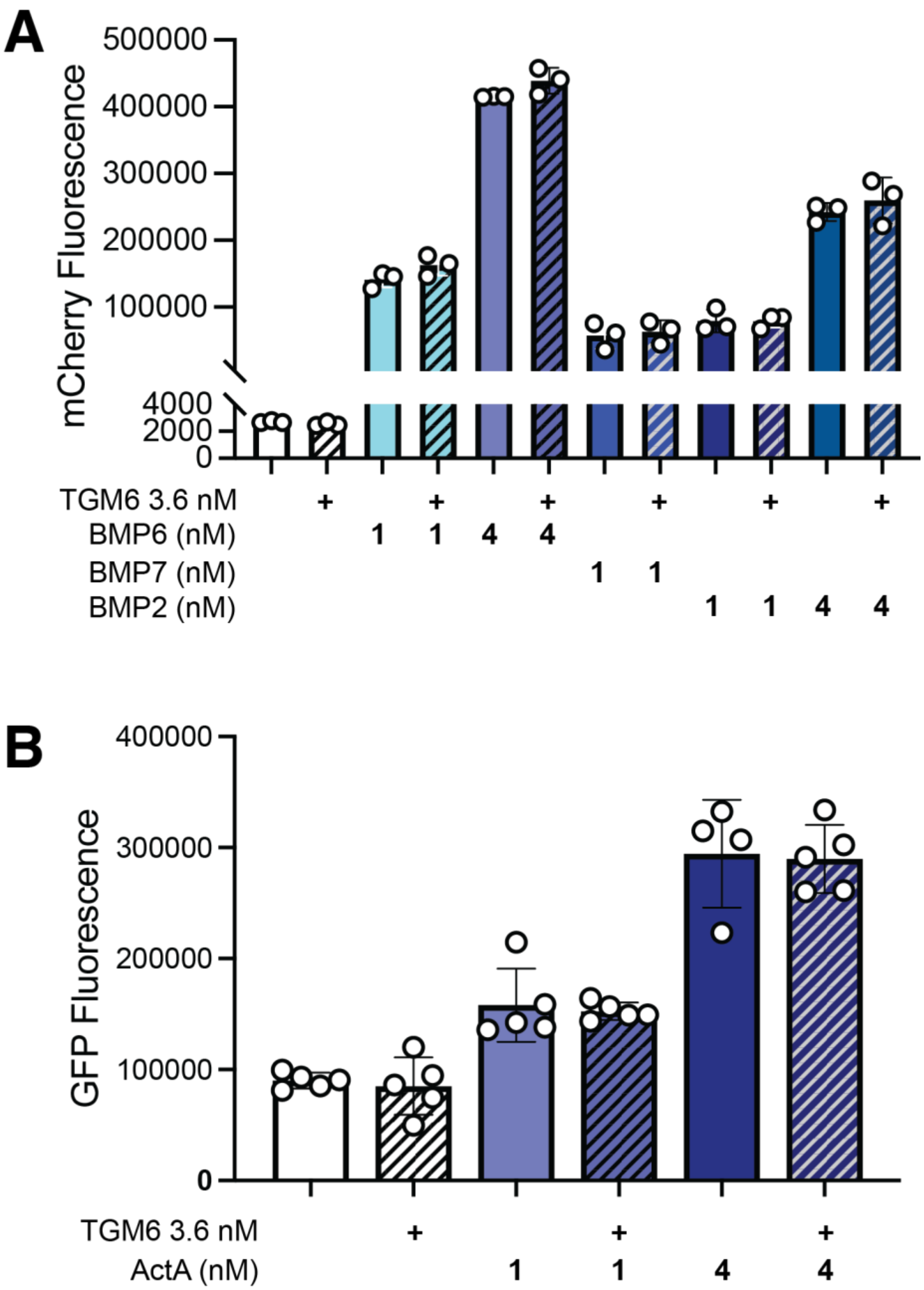

**Figure S7.** Overlay of the TβRII and TGM6-D3 components of the TβRII:TGM6-D3 crystal structure with crystal structure of unbound TβRII (PDB 1M9Z) or the lowest energy member of the ensemble of TGM1-D3 solution structures (PDB 7SXB), respectively.

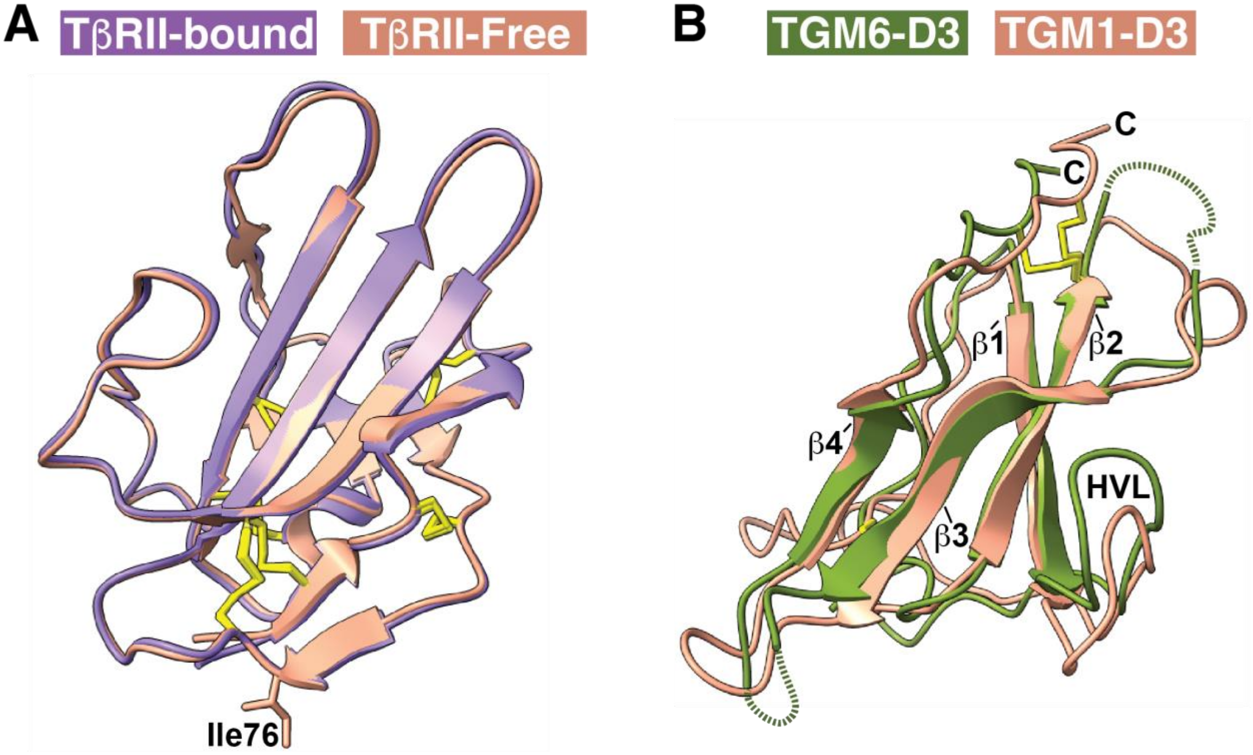

RMSD
Secondary Structure: 0.27 Å
Overall: 0.49 Å

RMSD
Secondary Structure: 0.96 Å
Overall: 1.25 Å

**Figure S8.** ITC thermograms obtained for WT TGM6-D3 injected into TβRII variants (A-E) and WT TβRII injected into TGM6-D3 variants (F-P) and TGM1-D3 variants (Q-T). (A) TβRII WT (B) TβRII D55A, (C) TβRII I76A, (D) TβRII D141A, (E) TβRII E142A, (F) TGM6-D3 WT, (G) TGM6-D3 R38A, (H) TGM6-D3 I78A, (I) TGM6-D3 Y80A, (J) TGM6-D3 Y80F, (K) TGM6-D3 R82A, (L) TGM6-D3 R82S, (M) TGM6-D3 Y93A, (N) TGM6-D3 R95A, (O) TGM6-D3 P94K R95N (KN), (P) TGM6-D3 Q81K R82S R83G G84T (KSGT), (Q) TGM1-D3 WT, (R) TGM1-D3 S242R, (S) TGM1-D3 K254P N255R (PR), and (T) TGM1-D3 K241Q S242R G243R T244G (QRRG).

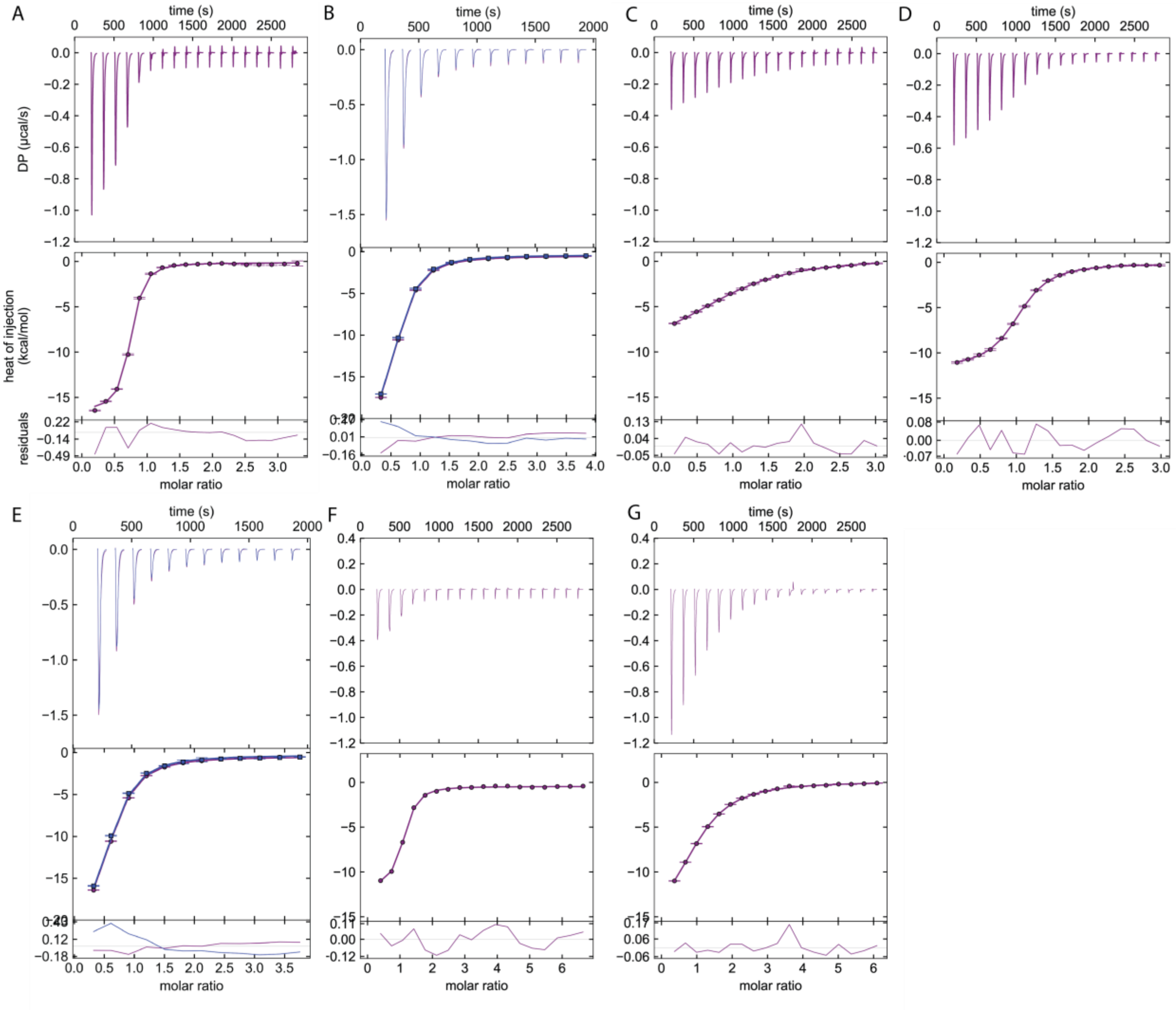

**Figure S8 continued**

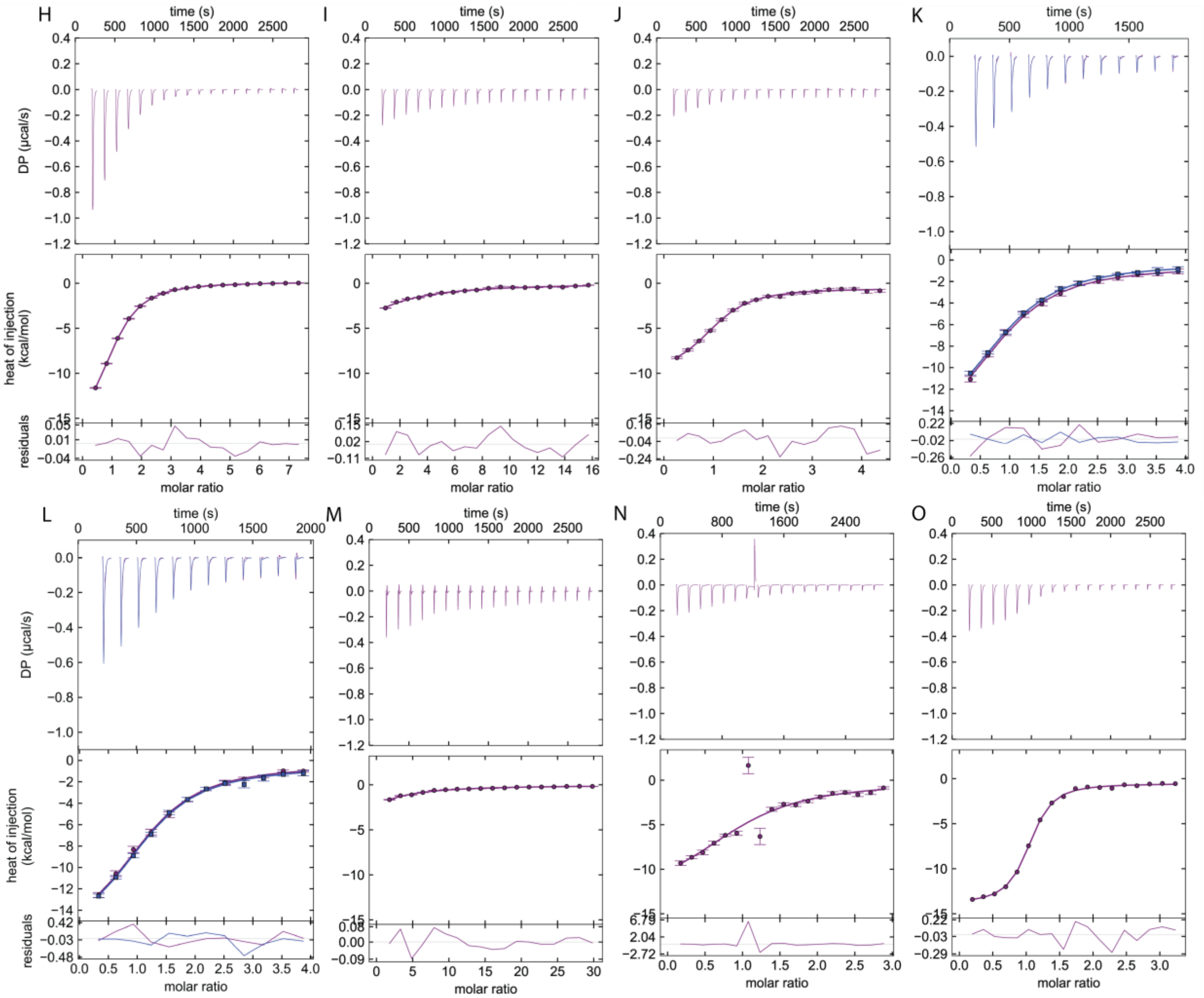

**Figure S8 continued**

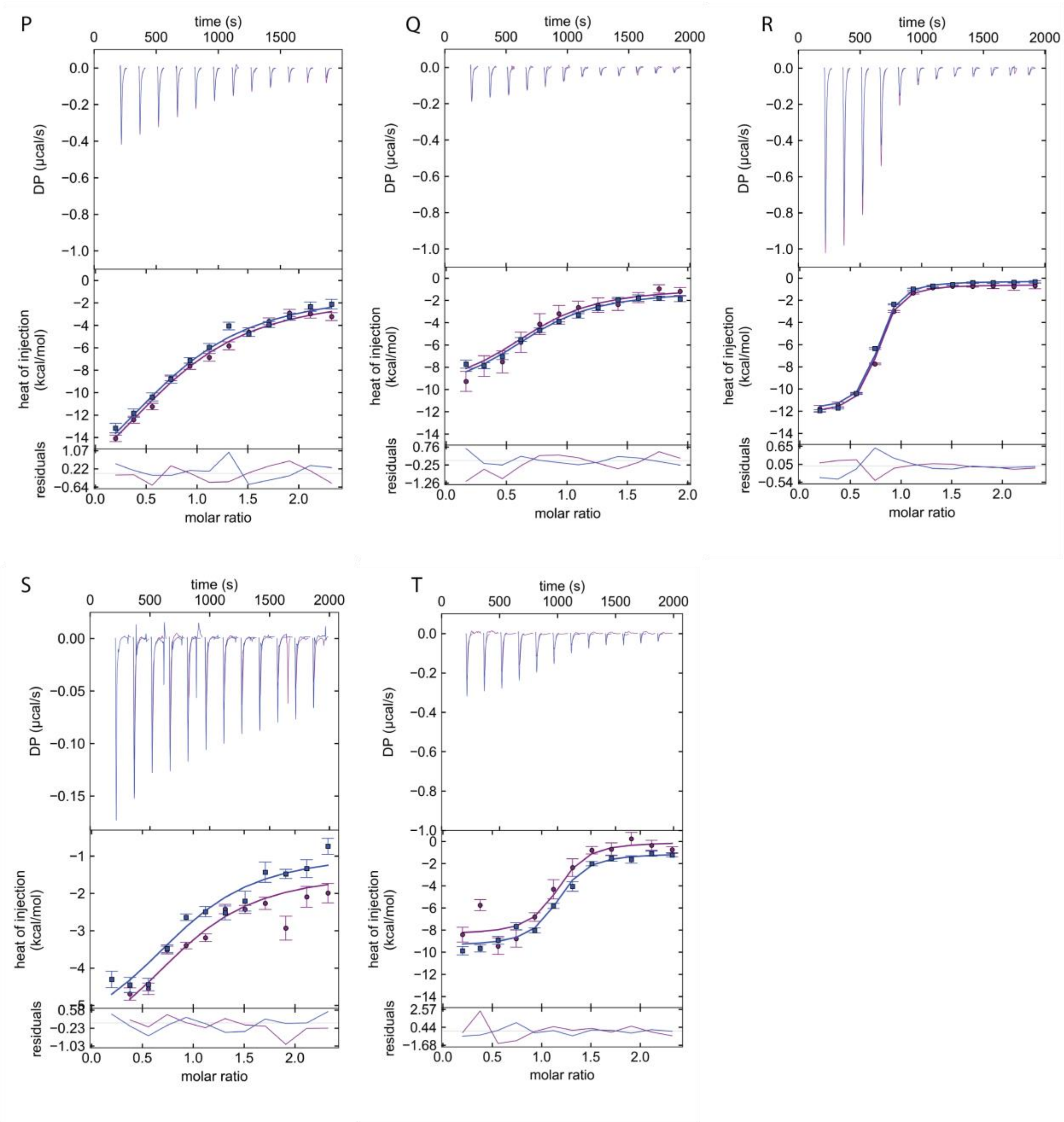

**Figure S9.** Methyl and amide regions of the 1D $^1H$ NMR spectra of the TGM6-D3 WT, I78A, Y80A, and Y93A variants used in the ITC experiments.

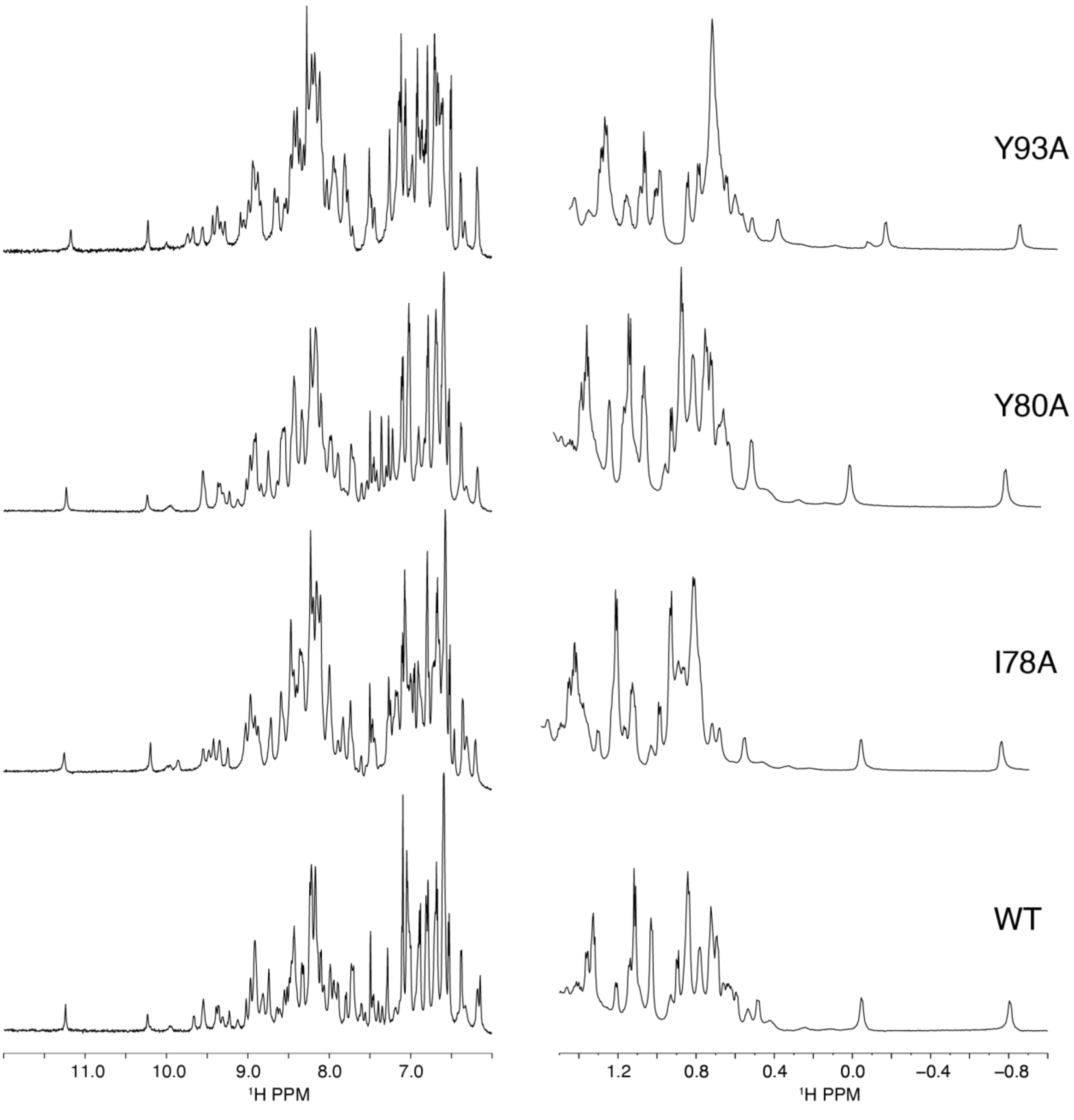

**Figure S10.** Omit map of TGM6-D3 residues 82-85 with model map at different contours: (A) 0.75σ, (B) 1.00σ, (C) 1.25σ, and (D) 1.50σ. Residues that were modeled in the final structure are displayed with transparency for reference but were not included during phasing.

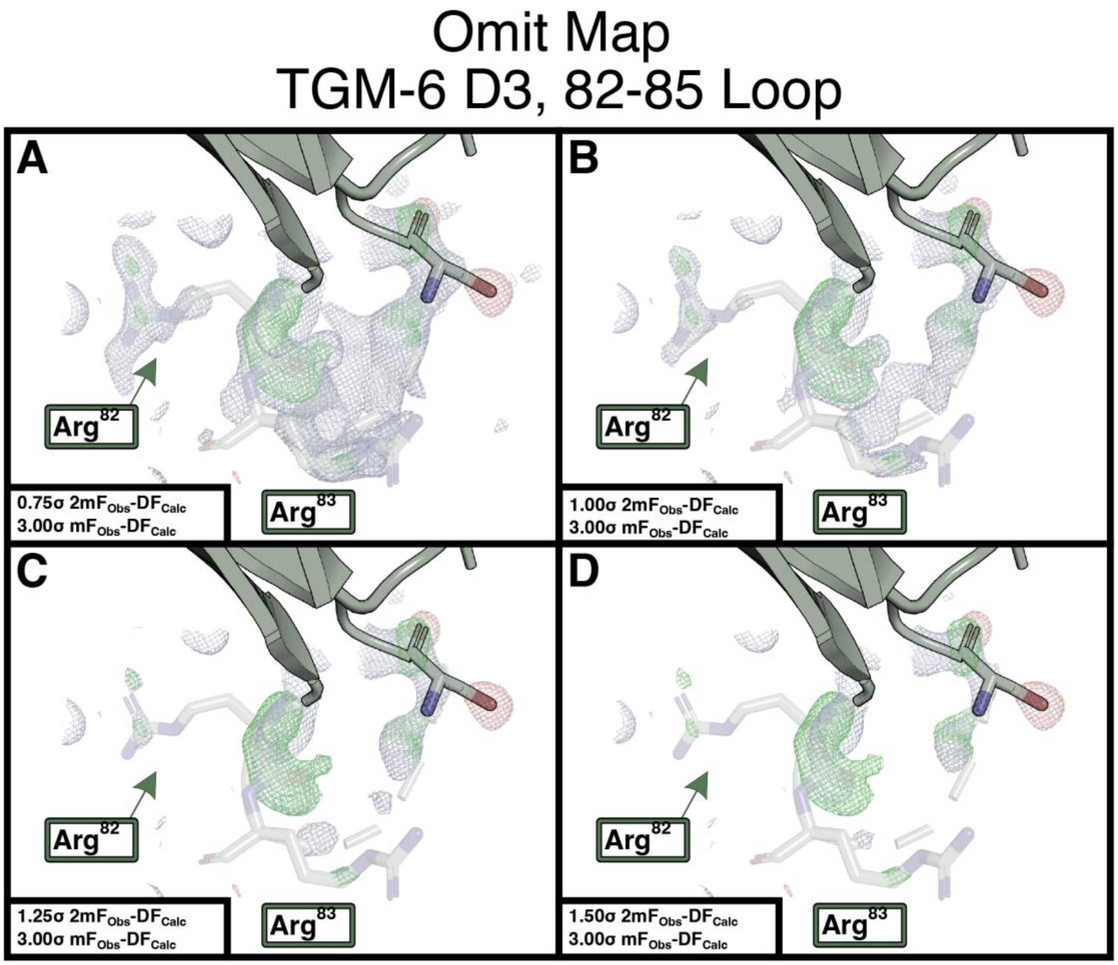

**Figure S11. D-F.** Inhibition of TGM1 signaling in MFB-F11 fibroblasts as detected by the conversion of p-nitrophenylphosphate by secreted alkaline phosphatase (**A-B**) or by pSMAD2 Western blotting (**C**) by either TGM6 (**A, C**) or TGM6 KSGT (**B, C**).

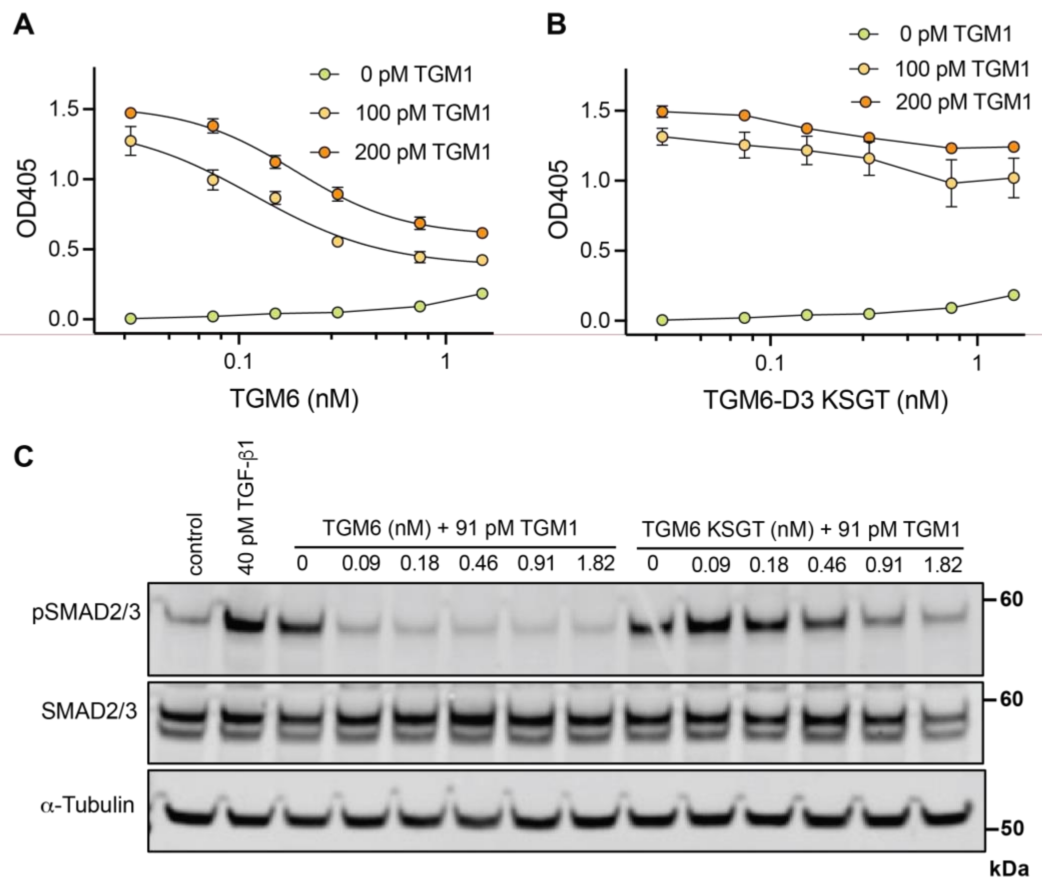

**Figure S12.** Alignment of the domain 3 amino acid sequences of TGM1-9**.** Conserved cysteines are highlighted in red. Residues highlighted in yellow correspond to those identified in TGM6 or TGM1 that contribute significantly to TβRII binding TβRII. Conservation of these residues in other TGMs is also indicated by yellow shading. TGM10 is not shown as this lacks domains 3 and 4 (Smyth, et. al, Int. J. Parisitology, 48, 379-385).

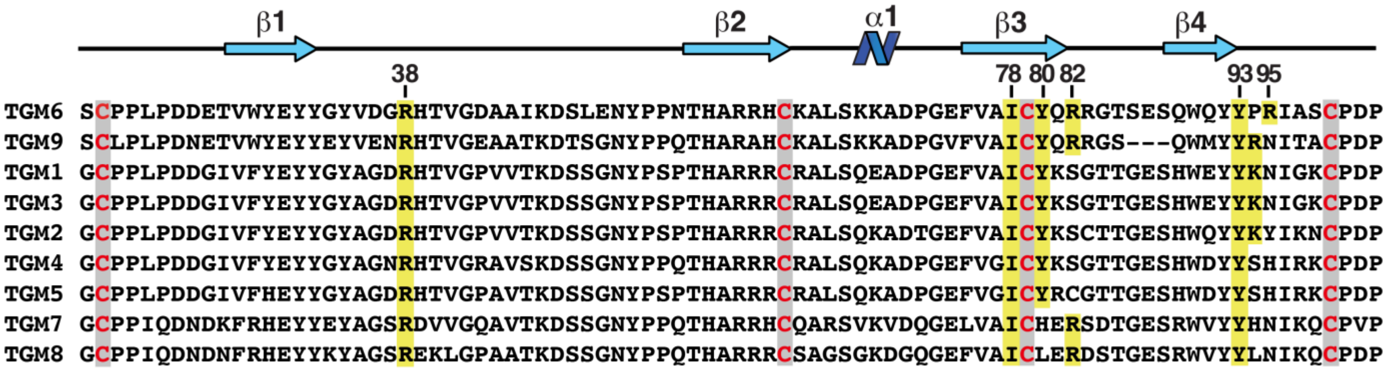
